# Contribution of interneuron subtype-specific GABAergic signalling to emergent sensory processing in somatosensory whisker barrel cortex in mouse

**DOI:** 10.1101/2021.02.18.431791

**Authors:** Liad J. Baruchin, Michael M. Kohl, Simon J.B Butt

## Abstract

Mammalian neocortex is important for conscious processing of sensory information. Fundamental to this function is balanced glutamatergic and GABAergic signalling. Yet little is known about how this interaction arises in the developing forebrain despite increasing insight into early GABAergic interneuron (IN) circuits. To further study this, we assessed the contribution of specific INs to the development of sensory processing in the mouse whisker barrel cortex. Specifically we explored the role of INs in speed coding and sensory adaptation. In wild-type animals, both speed processing and adaptation were present as early as the layer 4 critical period of plasticity, and showed refinement over the period leading to active whisking onset. We then conditionally silenced action-potential-dependent GABA release in either somatostatin (SST) or vasoactive intestinal peptide (VIP) INs. These genetic manipulations influenced both spontaneous and sensory-evoked activity in an age and layer-dependent manner. Silencing SST+ INs reduced early spontaneous activity and abolished facilitation in sensory adaptation observed in control pups. In contrast, VIP+ IN silencing had an effect towards the onset of active whisking. Silencing either IN subtype had no effect on speed coding. Our results reveal how these IN subtypes differentially contribute to early sensory processing over the first few postnatal weeks.

## Introduction

The mammalian neocortex is a higher order area of the central nervous system responsible for processing of sensory information and initiation of voluntary behaviour. Essential to this role are local circuits comprised of glutamatergic pyramidal cells and locally-projecting GABAergic interneurons (INs). These two populations integrate incoming sensory information – relayed via the thalamus – to generate percepts, which subsequently elicit an appropriate behavioural response through efferent pyramidal cells. Much of our understanding of the processes underpinning such computations – at the cellular and circuit level, has been derived from fundamental research in animal models. One such model is the mouse somatosensory barrel field (S1BF): the area of the neocortex responsible for processing incoming tactile sensory information arising from the whiskers (Petersen, 2007). Investigations performed in mature rodents have revealed that neurons in the columnar and layered structure of S1BF can derive various stimulus properties from incoming signals, such as location, speed, texture and relative novelty (Guić-Robles, Valdivieso, & Guajardo, 1989; Musall, Haiss, Weber, & von der Behrens, 2017; R. S. Petersen, Panzeri, & Diamond, 2002). A body of evidence has identified that GABAergic signalling (Rudy et al. 2011; Yu et al. 2019; Muñoz et al. 2017)) is required for such sensory processing in the mature neocortex (Ayzenshtat, Karnani, Jackson, & Yuste, 2016; Kolasinski et al., 2017; Natan et al., 2015; Wood, Blackwell, & Geffen, 2017). However, the contribution of GABAergic INs to nascent processing in the developing brain is still unknown.

In the developing neocortex there is an additional challenge, namely to balance emergent sensory processing and formative behavioural output with the need to integrate and establish circuit function. This challenge is met across primary sensory areas – including S1BF (Erzurumlu & Gaspar, 2012) – by gradual shifts in synaptic connectivity and plasticity, during which time there are changes in the nature of cortical activity. This includes oscillations not present in the adult neocortex such as the intermittent spontaneous spindle bursts (SB) (Rustom Khazipov et al., 2004; Marat Minlebaev, Ben-Ari, & Khazipov, 2007). To date, our understanding of which neuronal subtypes that contribute to these formative activity patterns and emergent perception is limited (Hanganu-Opatz et al., 2021). What is clear is that GABAergic interneuron diversity has a role to play in constraining the influence of early sensory input and sculpting early circuits (Butt, Stacey, Teramoto, & Vagnoni, 2017; Modol et al., 2020). Of the three main classes of interneuron (Rudy et al., 2011), parvalbumin (PV+)-expressing INs have been shown to play an important role in the closure of periods of plasticity (Hensch, 2005; McRae, Rocco, Kelly, Brumberg, & Matthews, 2007; Nowicka, Soulsby, Skangiel-Kramska, & Glazewski, 2009) and the onset of fast adult-like signalling (Doischer et al., 2008). In contrast, recent evidence has identified that one of the other prominent IN classes – defined by expression of the peptide somatostatin (SST+), contributes to early neurodevelopment events including synaptogenesis, sensory innervation and neuronal maturation(Marques-Smith et al., 2016; Oh, Lutzu, Castillo, & Kwon, 2016; Tuncdemir et al., 2016). The third main class are defined by expression of the ionotropic serotonin receptor, 5-HT_3A_R (Rudy et al., 2011). These are born late in embryonic development (S. Butt et al., 2005; Miyoshi et al., 2015) and as such are thought to contribute to circuit refinement towards the onset of active sensory perception (Hanganu-Opatz et al., 2021). That said, one major subtype of 5-HT_3A_R IN - the genetically-tractable vasoactive intestinal peptide-positive (VIP+) INs, have recently been shown to influence early circuits via their interaction with pyramidal cells and SST+ INs (Batista-Brito et al., 2017; Marques-Smith et al., 2016; Tuncdemir et al., 2016; Vagnoni et al., 2020). Based on this understanding, we hypothesised that both SST+ and VIP+ INs contribute to emergent sensory processing through postnatal life; a role that we can test using conditional silencing of neurotransmitter release via deletion of the SNARE complex protein Snap25 (Marques-Smith et al., 2016; Washbourne et al., 2002).

To assess the role that SST+ and VIP+ INs play in early cortical sensory computations, we recorded spontaneous activity and sensory-evoked responses from S1BF *in vivo* through the layer (L)4 critical period of plasticity (CP) up to, and including, the onset of active whisking (AW). We found that conditionally silencing SST+ IN signalling led to a reduction in spontaneous SBs during the CP in line with delayed thalamic innervation (Marques-Smith et al., 2016), whereas silencing VIP+ INs had no effect at this early age. At the onset of AW, silencing either IN subtype resulted in increased spike activity across the entire cortical column. In terms of sensory integration, we favoured multi-whisker as opposed to a single-whisker stimulation as this best captures the natural stimulus at early ages (Carvell & Simons, 1990; Kleinfeld, Ahissar, & Diamond, 2006). Beyond assessment of the simple sensory-evoked responses we also focused on speed coding and adaptation. These two processes, that have been previously studied around the onset of active sensation (van der Bourg et al., 2016), underlie more complex perceptual processing in the mature cortex (Allitt, Alwis, & Rajan, 2017; Arabzadeh, Petersen, & Diamond, 2003; Maravall, Petersen, Fairhall, Arabzadeh, & Diamond, 2007; Ollerenshaw, Zheng, Millard, Wang, & Stanley, 2014); perceptual processing that most likely requires temporal and spatial recruitment of diverse interneuron subtypes. We found that in wild-type animals speed was encoded in a consistent manner from the earliest time point tested. However, adaptation in the sensory response varied in profile over development. Silencing of GABAergic signalling in our two IN subtypes did not affect speed processing *per se*, but did result in altered sensory-evoked responses in an age and layer specific manner. This confirms differing roles for SST+ and VIP+ INs in emergent sensory processing in the developing somatosensory cortex.

## Methods

### Mouse Lines

Animal experiments were approved by the University of Oxford local ethical review committee and conducted in accordance with Home Office project (30/3052; P861F9BB7) licenses under the UK Animals (Scientific Procedures) 1986 Act. The following mouse lines, maintained on a mixed (C57B15/J || CD1) backgrounds were used: a conditional floxed-*Snap25* [Snap25<tm3mcw>] line and the VIP-ires-Cre [Vip<tm1(cre)Zjh>] and SST-ires-Cre [Sst<tm2.1(cre)Zjh>]. SST-ires-Cre^HOMO^;Snap25^C/+^ or VIP-ires-Cre^HOMO^;Snap25^C/+^ were crossed with Snap25^C/C^ mice to generate offspring with either functional (Snap25^C/+^) or silenced (Snap25^C/C^) SST+ or VIP+ INs respectively. All experiments were performed blind to the genotype, which was ascertained by PCR following completion of the data analysis.

### Surgical and recording procedures

Animals were anaesthetised with urethane (U2500; Sigma Ltd., UK) with a dose of 0.5-1g/kg. Depth of anaesthesia was verified by absence of reflexes and the animals’ breathing and heart rate were constantly monitored throughout the recording procedure thereafter. The animal was fixed to a stereotaxic frame (51600; Stoelting; UK) with a mouse adaptor (51615; Stoelting). Contralateral whiskers were fitted into a cannula attached to a piezo electric unit (Thor labs; PB4VB2W), connected to a piezoelectric amplifier (E-650 Piezo Amplifier; PI; Germany). The skull was then exposed and in animals younger than P10 was strengthened by applying a thin layer of cyanoacrylic glue (Loctite). Barrel cortex coordinates were identified using a neonatal brain atlas (Paxinos, Halliday, Watson, & Mustafa, 2020), and a small craniotomy was made with a surgical drill (Volvere i7, NSK Gx35EM-B OBJ30013 and NSK VR-RB OBJ10007). A silicon probe (Neuronexus A1×32-Poly2-5mm-50s-177-A32) covered in DiI solution (1,1’-dioctadecyl-3,3,3’,3’-tetramethyl indocarbocyanine; Invitrogen; UK) was implanted by lowering it slowly into the brain. After a minimum of 30 minutes post-implantation, a baseline period of 20 minutes was recorded, after which the experimental protocols were conducted.

### Experimental procedures

#### Whisking speed manipulation

In each trial, a single whisker deflection was delivered at varying velocities with rectified sine waves with a width equivalents of 5, 10 20, 40 and 80 Hz. The interval between deflections was 30 seconds. The different speeds were given in consecutive blocks of 20 trials, with a different order for each mouse.

#### Paired-pulse procedure

In each trial two consecutive whisker deflections of 80 Hz/288 deg/ms were delivered with a varying inter-stimulus interval of 100, 250, 500, 1000 and 1500 ms. The inter-trial interval was 30 second, and each ISI was delivered in consecutive blocks of 20 trials, with different order for each mouse.

### Current-Source Density analysis and Layer Localisation

The current-source density (CSD) maps were derived using a previously published method (Nicholson & Freeman, 1975) with a correction for the topmost and bottommost electrodes suggested by Vaknin (Vaknin, DiScenna, & Teyler, 1988). The estimated CSD, C at depth z is described as:

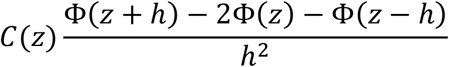

Where *ϕ* is the potential at a specific depth and h is the vertical spacing between the electrodes. Each pair of contacts was averaged when using the procedure to increase the signal to noise ratio. To reduce spatial noise further, we applied the three-point hamming filter (Rappelsberger, Pockberger, & Petsche, 1981):

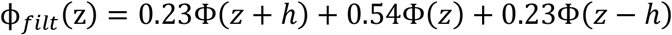

The shortest latency, large amplitude sink was classified as the granular layer, the contacts above as the supragranular layers, and those below it as the infragranular layers (M. Minlebaev, Colonnese, Tsintsadze, Sirota, & Khazipov, 2011; Nicholson & Freeman, 1975). In each layer, only contacts that showed consistent activity and a 50 Hz noise below 2 standard devations (SDs) of the power spectrum curve were chosen for further analysis, while the same number of chosen contacts was used for analysis for each of the layers.

### Data Analysis

Data analysis was performed post hoc in Matlab (Matlab 2019b).

#### Baseline activity

in order to identify spindle burst (SB) activity, we first filtered the signal between 5 and 35 Hz and then isolated periods where the signal exceeded 2 SD of the average signal. Events with a duration of less than 100 ms and/or less than 3 cycles were discarded. The frequency of the SBs was calculated as the sum of troughs divided by the duration of the event. The baseline firing rate was taken by calculating using a running average of 500 ms and then averaging the windows together to get the gross-average.

#### Spike Sorting

spike sorting was performed using Kilosort2 (Pachitariu, Steinmetz, Kadir, Carandini, & Kenneth D., 2016; github.com/cortex-lab/KiloSort). After the automatic classification of spikes into units, manual verification was performed using phy (github.com/kwikteam/phy). These spikes were then combined into a multi-unit signal for each layer.

#### Sensory-evoked response (SER)

for both MUA and LFP, the evoked sensory response was derived by averaging across trials, and then finding the peak of the resulting deflection. This was defined as the response amplitude (mV or Hz accordingly), and the time from the onset of whisker deflection to this point was defined as the peak latency (ms). This procedure was repeated for each of the identified layers. To correct for differences in baseline, a baseline subtraction was performed using a baseline of 100 ms before the whisker deflection.

#### Paired-pulse ratio (PPR)

the PPR was calculated by dividing the peak amplitude of the second response by the peak amplitude of the first response.

### Histology

At the end of the experiments, following administration of terminal anaesthesia, the brains were dissected and immersed in 4% paraformaldehyde (PFA; Alfa Aesar) in phosphate-buffered solution (PBS, Sigma) for 2 days. The brain was washed three times in a PBS solution, and cut into 80µm thick coronal slices using a vibratome (Leica VT1000S). To assist barrel localization, slices were counterstained with 4’,6-diamidine-2’-phenylindole 502 dihydrochloride (DAPI, D3571, Molecular Probes; dilution 1:1000) for 5 minutes followed by 2 minutes wash in PBS. Slices were mounted on a slide and imaged using either widefield fluorescent or confocal microscopy (LSM710; Zeiss) to verify the location of the electrode.

### Statistical Analysis

Statistical analysis was performed using Prism (Graphpad). Normality of the data was tested using the Shapiro-Wilk test. Differences in populations with normal distributions were tested using Student’s t test or one-way ANOVA. In cases where normality assumptions were violated, Mann-Whitney (M-W), Kruskal-Wallis (K-W), and Wilcoxon tests were used. Bonferroni correction (BfC) and Dunn test (Dunn) were applied for multiple comparisons as appropriate. Alpha levels of p ≤ 0.05 were considered significant. Data are presented as the mean ± standard error of the mean (SEM).

## Results

### The Development of Sensory-Evoked Responses in mouse S1BF

We performed multi-electrode *in vivo* electrophysiology to record both local field potential (LFP) and multi-unit activity (MUA) across the depth of S1BF in response to multi-whisker stimulation across early development. We divided our analysis into three developmental time windows: postnatal day (P)5-P8 (equivalent to the layer 4 critical period for plasticity; CP), the next few days prior to active perception (P9-P11; pre-active whisking; pre-AW) and a time window covering the onset of active perception (P12-P16; active whisking; AW); windows that reflect our existing knowledge of underlying circuitry from *in vitro* studies (Anastasiades & Butt, 2012; Anastasiades et al., 2016; Vagnoni et al., 2020). Our initial analysis focused on granular layer 4 (G) as this layer showed robust LFP and MUA across all three developmental time points (**Figure 1A**,**B**).

**Figure 1.**
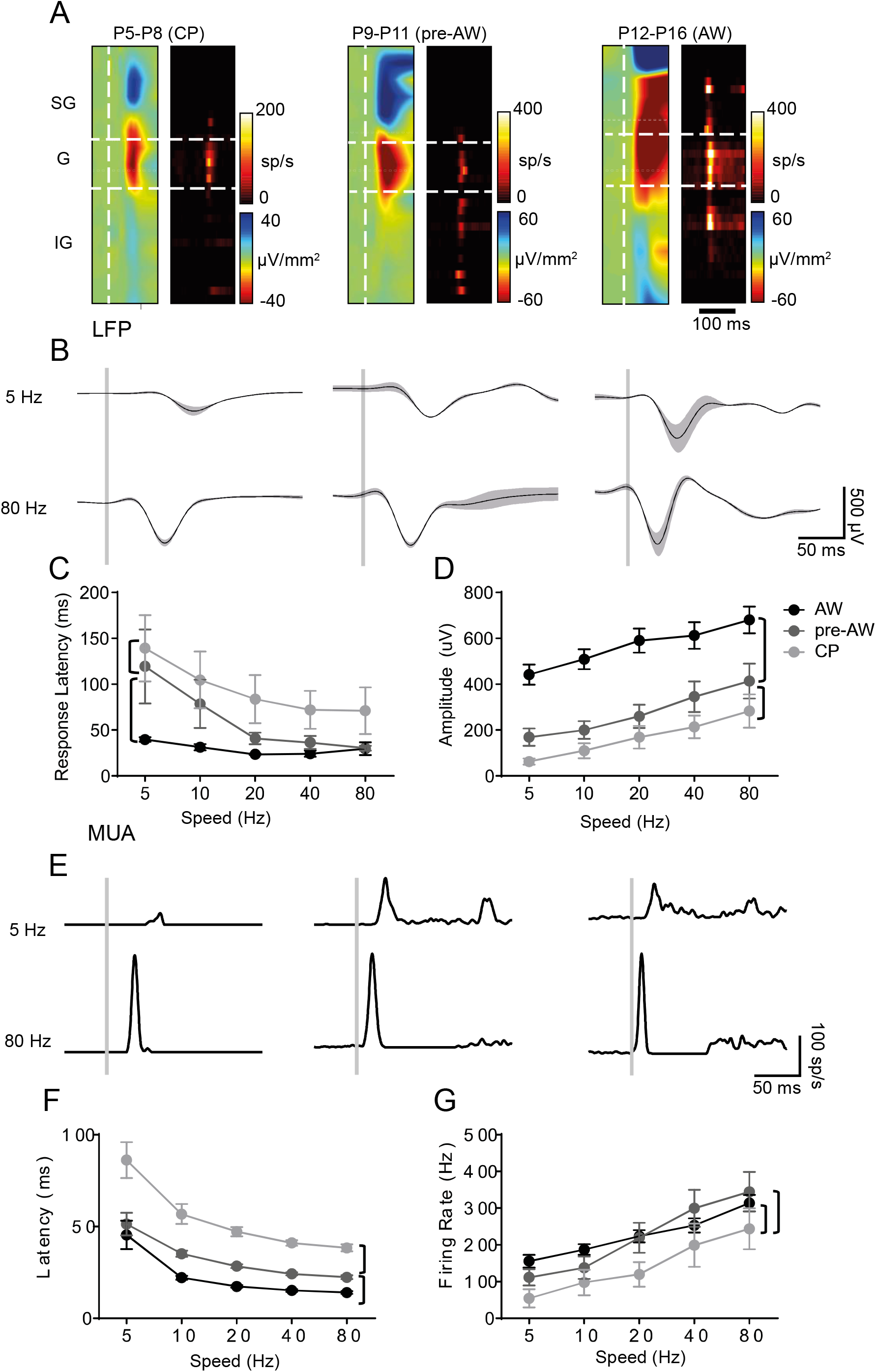
Sensory evoked response (SER) amplitude increases through development. **A**. CSD and MUA plots across the depth of the cortex after a single multi-whisker deflection (indicated by vertical white dashed line), during CP (P5-P8), pre-AW (P9-P13) and AW (P14-P21); SG, supragranular; G, granular; IG, infragranular layers. **B**. Granular layer LFP responses in the aforementioned time periods after a 5Hz or 80 Hz whisker deflection (onset indicated by vertical grey bar). **C**. Averages of the LFP response latencies for the different deflection speeds across the developmental time periods. There was a significant effect for both age (F (2, 188) = 22.19, p < 0.01) and speed (F (4, 188) = 5.801, P <0.01). **D**. Averages of the LFP response amplitudes for the different deflection speeds across the developmental time periods. There was a significant effect for age (F (2, 188) = 75.92, p < 0.01) and speed (F (4, 188) = 7.135, p<0.01). **E**. Typical granular layer MUA response in the aforementioned time periods after a 5Hz or 80 Hz whisker deflection. **F**. Average MUA response latencies across the developmental time-periods. There was a significant change with speed (F(4,159) = 25.27, p<0.01) and with age (F(2,159) =63.59, p<0.01). **G**. Average MUA response peak firing rates across the developmental time-periods. There was a significant change with speed (F(4,159) = 17.51, p<0.01) and with age (F(2,159) =11.16, p<0.01). LFP: CP: N=11, pre-AW: N = 9, AW: N=21, MUA: CP: N = 8, pre-AW: N = 6, AW: N = 21. Brackets: p<0.05 difference in an ANOVA multiple comparison.

We found that the amplitude and latency of the LFP evoked response scaled with deflection speed starting as early as CP (**Figure 1B-D**), indicative of differentially encoding of whisking speed prior to active perception (AW). The decrease in latency (**Figure 1B**,**C**) and increase in amplitude (**Figure 1B**,**D;** p<0.01) of the LFP across developmental time windows suggests that an increase in thalamic synaptic input – most prominent between pre-AW and AW time windows – could be a major influence on the observed sensory responses (Malina, Mohar, Rappaport, & Lampl, 2016).To further examine any cortical contribution to early encoding of speed we further examined MUA responses to different deflection speeds (**Figure 1E**). Similar to the LFP, the response latency and amplitude were sensitive to speed within developmental time windows (**Figure 1E-G**). However, the frequency of the sensory response reached a maximum during pre-AW (p<0.01). The LFP in granular layer is a sum of both thalamic synaptic input and intra-cortical activity (Malina et al., 2016), this suggests that maturing intra-cortical inhibition (Daw, Ashby, & Isaac, 2007) counterbalances the increasing thalamic signalling.

To study the development of adaptation we used a paired-pulse paradigm – two identical stimuli presented in sequence with varying inter-stimulus intervals (ISIs), similar to previous research (van der Bourg et al., 2019, Zehendner et al., 2013), however, over a larger range of developmental time points (CP to AW). We recorded LFP and MUA (**Figure 2A**) in response to a paired-pulse paradigm of varying ISI (between 0.1 and 1.5s), focusing our analysis on granular layer 4 (**Figure 2B-E**). Prior to AW, we observed failure of the second response at the shortest ISI test (0.1s) in 12 of 22 animals during CP and 13 out of 25 during pre-AW. No such failure was observed during AW. In the older animals, analysis of LFP (**Figure 2B**) showed a pattern of depression that becomes weaker with longer ISIs. However, during the CP time window, the paired-pulse ratio (PPR) profile takes a distinctive ‘reverse-U’ shape in respect to ISI (**Figure 2D**), whereby the PPR peaked at 0.5s and fell away at shorter (0.1s: p<0.01 compared to AW) and longer ISIs (1.5s; p<0.01 compared to AW). The same was true during pre-AW, although to a lesser degree (p<0.01 for 0.1s and 1.5s compared to AW). Analysis of MUA (**Figure 2C**,**E**) revealed a similar pattern in PPR over varying ISI, with the exception of.0.5s ISI during the CP (Figure 2E). Stimulating whiskers at 0.5s ISI within CP resulted in a larger MUA in response to the second stimulation (p<0.01 compared to pre-AW and AW) and the only positive PPR observed in L4 MUA across early development.

**Figure 2.**
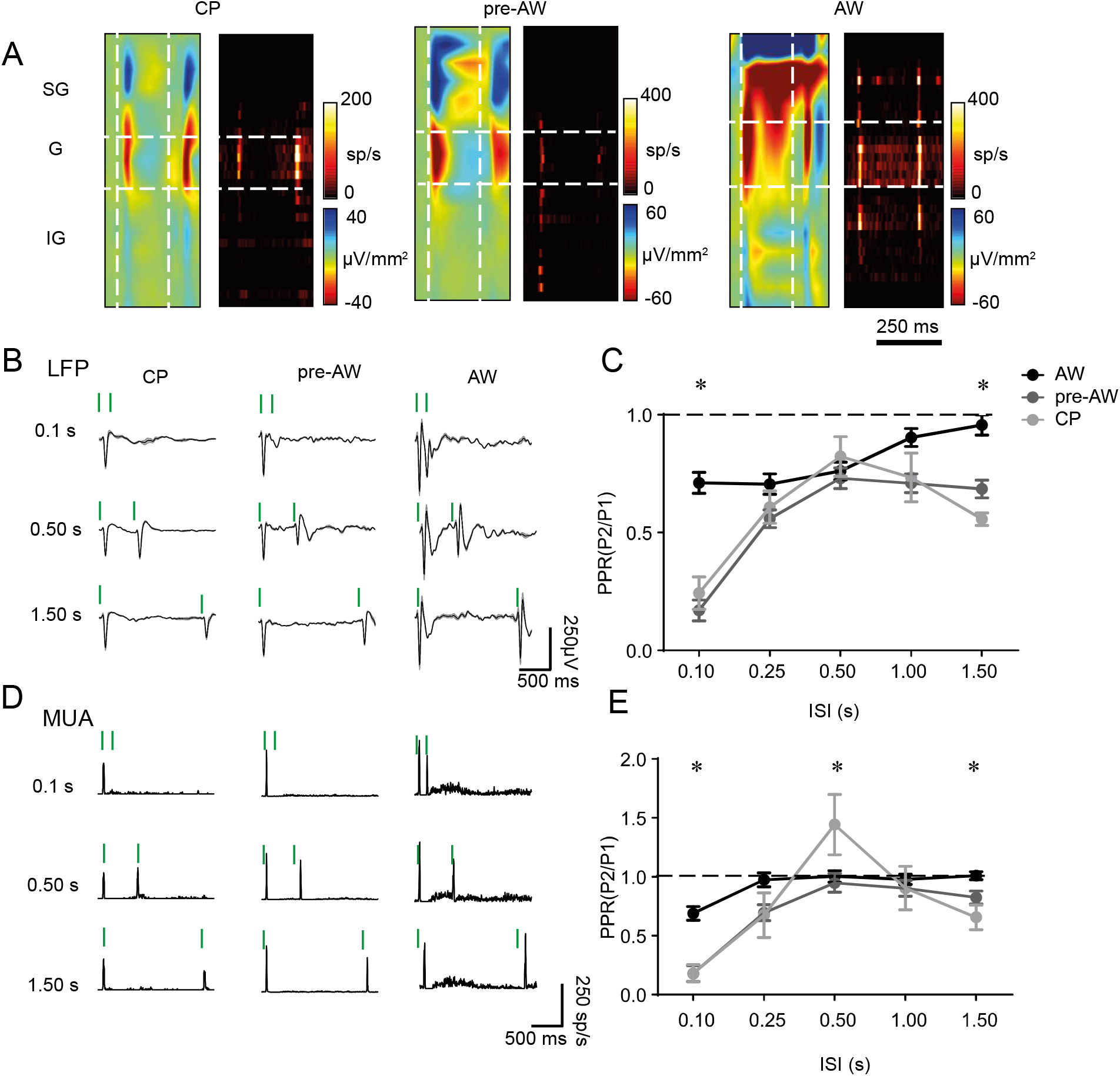
Adaptation pattern changes through development. **A**. CSD and MUA plots showing the sensory response across the depth of the cortex after two deflections with a 0.25s ISI, during CP, pre-AW, and AW. **B**. Typical granular layer LFP responses during the different time-periods for two deflections with ISIs of 0.1s, 0.50s, and 1.50s. **C**. Average PPR of the LFP response in the granular layer during CP, pre-AW, AW for ISIs of 0.10s, 0.25s, 0.50s, 1.00s, and 1.50s. There was a significant Age X ISI interaction (F (8, 307) = 3.272, p<0.001; CP: N = 22, pre-AW: N = 25, AW: N = 20). **D**. Typical granular layer MUA responses during the different time-periods for 2 deflections with ISIs of 0.1s, 0.50s, and 1.50s. **E**. Average PPR of the MUA response in the granular layer during CP, pre-AW, AW for ISIs of 0.10s, 0.25s, 0.50s, 1.00s, and 1.50s. There was a significant age X ISI interaction (F (8, 198) = 3.876, p<0.001; CP: N = 12, pre-AW: N = 16, AW: N = 18). Green bars show deflection times. * p<0.05 in a simple multiple comparison after an ANOVA.

### Impact of silenced interneuron signalling on spontaneous cortical activity in neonatal S1BF

GABAergic INs play an important role in cortical circuit formation and maturation (Ben-Ari, Khalilov, Represa, & Gozlan, 2004; S. J. Butt et al., 2017; Modol et al., 2020). However there is little understanding of how GABAergic IN diversity contributes to early activity on the millisecond time scale despite the observation that application of GABA antagonists can disrupt whisker-evoked activity (M. Minlebaev et al., 2011). To address the role for GABAergic IN signalling, we used a conditional *Snap25* knockout line (*Snap25*^*C/C*^), to produce mutant animals in which we conditionally abolished action potential-dependent release of GABA (‘silenced’) in SST+ (*SSTCre;Snap25*^*C/C*^; termed SST^cs^) or VIP+ (*VIPCre;Snap25*^*C/C*^; VIP^cs^) INs (Marques-Smith et al., 2016; Washbourne et al., 2002), and compared them to wild-type (WT) animals. We first investigated the impact of conditional silencing of SST+ or VIP+ INs on spontaneous spindle bursts (SB), synchronous neural activity, that is observed in early development, equivalent to our CP window (Rustom Khazipov et al., 2004; Marat Minlebaev et al., 2007). Across all three genetic backgrounds we observed oscillatory network events in the absence of sensory input (**Figure 3A**),and quantified their oscillation frequency and duration (**Figure 3B**). SBs in WT pups had similar rate of occurrence, duration and frequency (**Figure 3C-E**) to previous studies in mouse (Khazipov & Milh, 2017; Seelke & Blumberg, 2010). However, we observed a decrease in SB occurrence in SST^cs^ animals when compared to both WTs (p<0.01) and VIP^cs^ (p<0.01) animals (**Figure 3C**). We observed no difference in the duration (**Figure 3D**) and intra-spindle frequency (**Figure 3E**) of SB events across all 3 backgrounds. These findings are consistent with the role of thalamus on SB generation (Khazipov et al., 2004). The reduced SB occurrence in SST^cs^ pups can explained in part by weaker thalamic innervation of L4 spiny stellate neurons reported in SST^cs^ animals *in vitro* (Marques-Smith et al., 2016).

**Figure 3.**
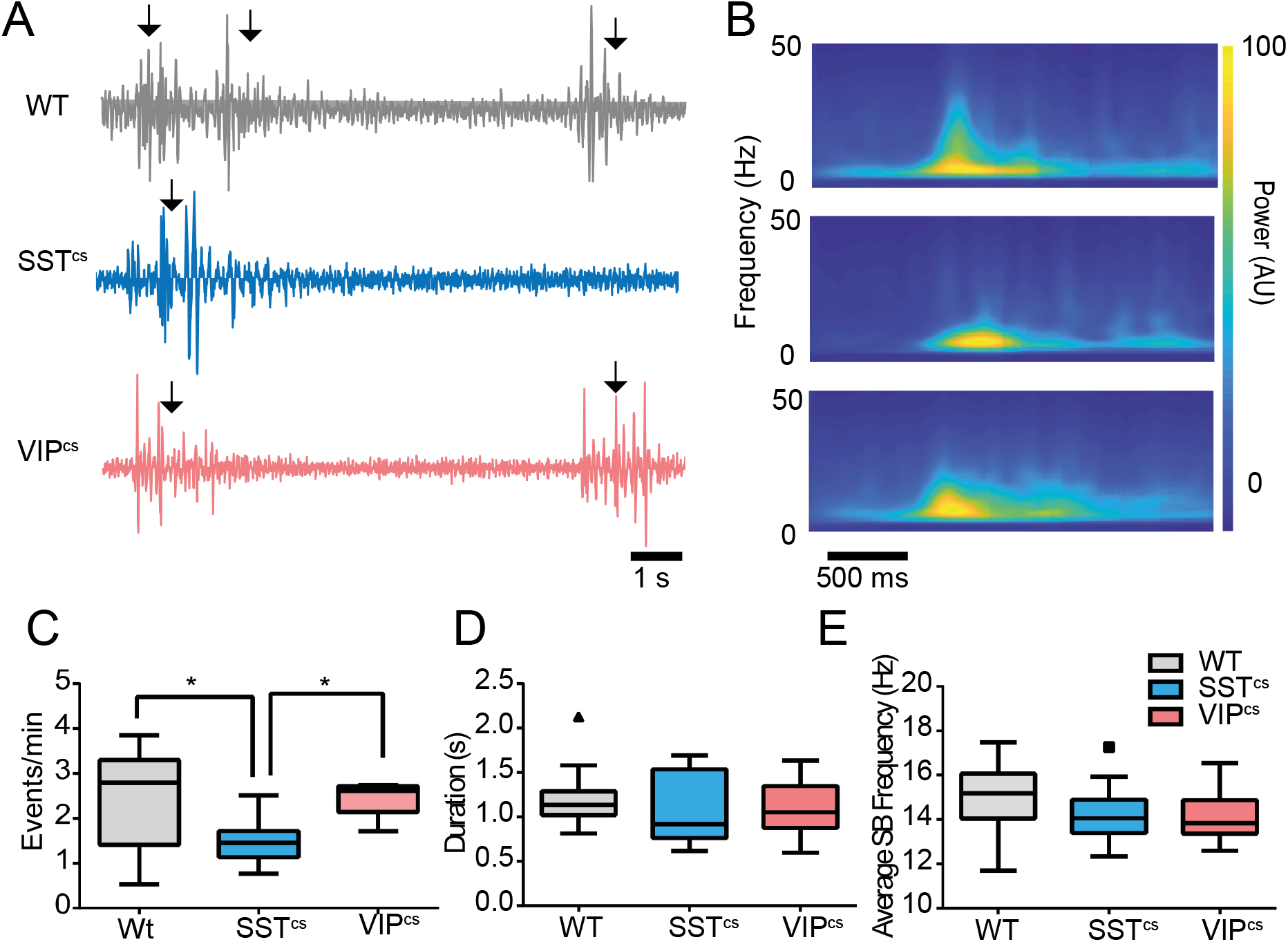
Silencing SST^cs^ leads to a decrease in SBs. **A**. Typical spontaneous activity during CP in WT, SST^cs^ and VIP^cs^ animals. SBs are marked by arrows. **B**. Average spectrograms from the recorded SBs in the 3 genetic lines. **C**. Average occurrence rate of SBs in WTs, SST^cs^ and VIP^cs^ animals. There was a significant difference between genotypes (F (2, 43) = 6.177, p <0.01). **D**. Average duration of the recorded SBs. There was no difference by genotype (F (2, 43) = 0.6287, P = 0.53). **E**. The average oscillations frequency of the recorded SBs. There was no difference by genotype (F (2, 43) = 1.533, p = 0.23). WT: N = 24, SST^cs^: N = 10, VIP^cs^: N = 12. *p<0.05 in post-hoc multiple comparisons.

We next examined the impact of silencing these INs subtypes on spontaneous cortical action potential discharge (MUA) both during CP and the later time windows (**Figure 4A**,**B**). We found that cortical firing is reduced in SST^cs^ animals during CP compared to WT (p<0.05), while VIP^cs^ animals had comparable cortical firing to WT (**Figure 4C**). During the pre-AW time window, we observed no difference in the spontaneous firing rate across all three background (**Figure 4D**). At the onset of AW there was a considerable increase in spontaneous firing rate (**Figure 4B**,**E**). This was more pronounced in both SST^cs^ (p<0.01) and VIP^cs^ (p<0.01) animals which had a higher firing rate than WT (**Figure 4B**,**E**). This observation is consistent with SST+ and VIP+ INs regulating spontaneous cortical activity at this later age. The reduced spiking during CP in SST^cs^ pups most likely reflects the reduction in SBs, since cortical firing is mostly limited to these events during this period (Rustom Khazipov et al., 2004). This further supports a role for SST+ INs in early network formation and function (Marques-Smith et al., 2016; Tuncdemir et al., 2016), one that later switches to an adult-like suppression of cortical activity (Urban-Ciecko, Fanselow, & Barth, 2015).

**Figure 4.**
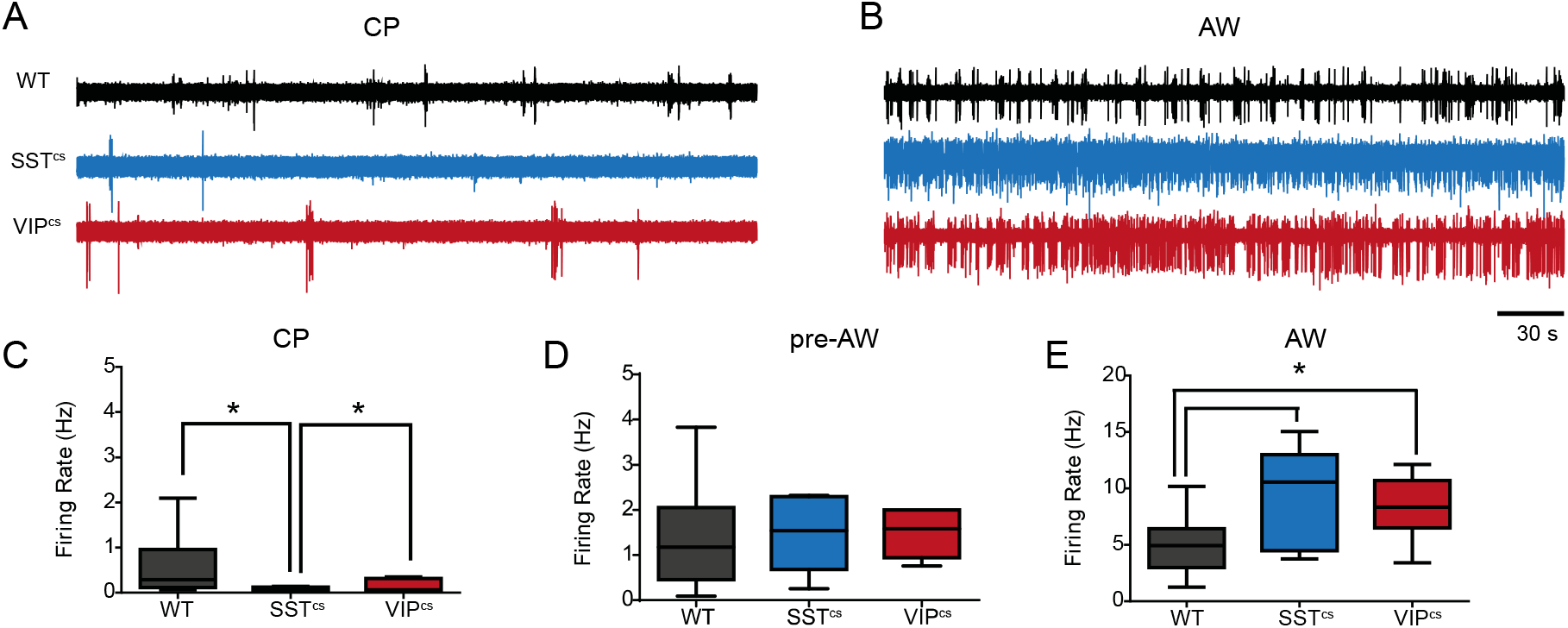
SST+ and VIP+ INs control spontaneous action potential discharge in an age dependent manner. **A**. Typical spontaneous spiking activity in WT, SST^cs^ and VIP^cs^ animals during CP. **B**. Typical spiking activity of these animals during AW. **C**. The average firing rate in WT, SST^cs^, and VIP^cs^ animals during CP. There was a significant different between genotypes (F(2,21) = 3.956, p<0.05; WT: N = 12, SST^CS^: N=7, VIP^cs^: N = 5). **D**. The average firing rate in these genotypes during pre-AW. There was no significant effect for genotype (F(2,21) = 0.048, p= 0.952. WT: N = 13, SST^CS^: N = 5, VIP^cs^: N = 6) **E**. The average firing rate in these genotypes during AW. There was a significant effect for genotype (F(2,31) = 6.850, p<0.01. WT: N = 22, SSTcs: N = 6, VIPCS: N = 6). * p<0.05 in a multiple comparison after an ANOVA.

### Contrasting Dynamics of SST+ and VIP+ in Sensory Responses

Our observation that INs differentially modulate spontaneous activity through development led us to ask how these interneurons might contribute to emergent sensory processing. We first examined how IN silencing affects the response to a single multi-whisker stimulation presented at different speeds. As before (**Figure 1**), we focused on our analysis on the LFP in the granular layer (**Figure 5A-C**). Silencing SST+ or VIP+ IN populations had no effect on the ability to encode speed across early development: across all three backgrounds and development time windows, we consistently observed a decrease in peak latency (**Figure S1A-C**) and an increase in amplitude (**Figure 5A-C**) for the L4 sensory responses according to speed. However, we observed an effect of IN silencing on the amplitude of the neural response to a single stimulus. During CP, both SST^cs^ (p<0.05) and VIP^cs^(p<0.01) animals exhibited a reduced amplitude of the sensory response compared to controls (**Figure 5A)**. During pre-AW, the response amplitude observed in SST^cs^ animals was comparable to WT, whereas VIP^cs^ animals exhibited a larger response than both WTs and SST^cs^ (**Figure 5B**; p<0.01 in both cases). This changed during AW: VIP^cs^ animals were indistinguishable from WT, whereas SST^cs^ animals exhibited delayed latency (**Figure S1C;** p<0.01 compared to WTs and VIP^cs^) and reduced amplitude of the sensory response compared to both WTs and VIP^cs^ animals (**Figure 5C**; p<0.01 compared to WT and VIP^cs^). Analysis of MUA (**Figure 5D-F**) revealed a similar pattern to that observed in the LFP with one significant exception: During AW, MUA in L4 of SST^cs^ was indistinguishable from that recorded in WT and VIP^cs^ animals (**Figure 5F**), consistent with delayed maturation of thalamic input into L4 S1BF in these animals (Marques-Smith et al., 2016)

**Figure 5.**
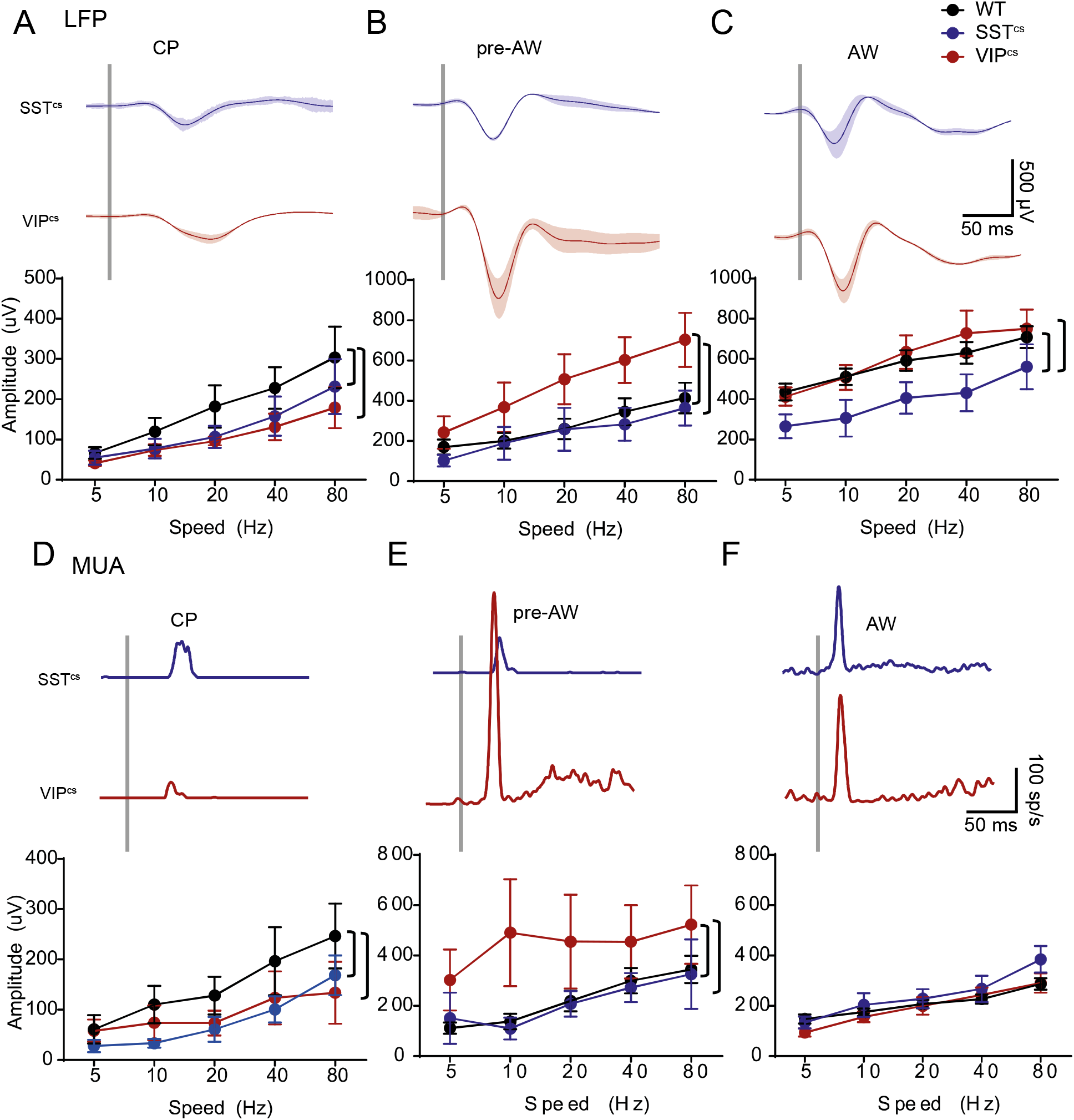
Time-dependent effect of SST+ and VIP+ IN silencing on SER. **A**. Top: LFP response after an 80Hz deflection in the granular layer of SST^cs^ and VIP^cs^ animals during CP. Bottom: the average peak amplitude of WT (n = 11), SST^cs^ (n=7) and VIP^cs^ (n=6) animals during CP. There was a significant effect for speed (F(4,98) = 6.743, p<0.001) and genotype (F(2,98) = 3.665, p < 0.05). **B**. Top: LFP response in the granular layer of SST^cs^and VIP^cs^ animals during pre-AW. Bottom: the average peak amplitude of WT (n=9), SST^cs^ (n=5) and VIP^cs^ (n=6) animals during pre-AW. There was a significant effects for speed (F(4,85) = 6.737, p<0.001) and genotype(F(2,85) = 11.59, p<0.001). **C**. Top: LFP response in the granular layer of SST^cs^ and VIP^cs^ animals during AW. Bottom: average peak amplitude of WT (n=21), SST^cs^ (n=6) and VIP^cs^ (n=6) animals during AW. There was a significant effects for speed (F(4,152) = 7.205, p<0.001) but not for genotype (F(2,152) = 9.600, p<0.001). **D**. Top: MUA response after an 80Hz deflection in the granular layer of SST^cs^ and VIP^cs^ animals during CP. Bottom: average peak firing-rate of WT (n=8), SST^cs^ (n=5) and VIP^cs^(n=5) animals during CP. There was an effect for speed (F(4,70) = 4.58, p<0.001), and genotype (F(2,70) = 3.834, p <0.05).**E**. Top: MUA response in the granular layer of SST^cs^and VIP^cs^ animals during pre-AW. Bottom: average peak firing-rate of WT (n=6), SST^cs^(n=4) and VIP^cs^ (n=6) animals during pre-AW. There was no effect for speed (4,60) = 1.650, p = 0.0.173), but there was an effect for genotype (F(2,60) = 7.208, p<0.01). **F**. Top: MUA response in the granular layer of SST^cs^ and VIP^cs^ animals during AW. Bottom: average peak firing-rate of WT (n=21), SST^cs^ (n=6) and VIP^cs^ (n=6) animals during AW. There was an effect for speed (F(4,147) = 16.81, p<0.001), but not for genotype (F(2,147) = 2.404, p = 0.094. Brackets show a p<0.05 multiple comparison after an ANOVA.

### A role for SST+ but not VIP+ INs in paired-pulse facilitation

We examined the paired-pulse response in the LFP (**Figure 6A**,**B**) and found that in both SST^cs^ and VIP^cs^ there was a linear relationship between PPR and ISI. PPR increased in SST^cs^ across the developmental windows tested (**Figure 6C**; p<0.01 between all age groups) whereas in VIP^cs^ animals there was a decrease in PPR during pre-AW not observed in WT and SST^cs^ animals (**Figure 6D**; p<0.01 compared to both CP and AW). We then compared the PPR between the three backgrounds across the different time-points. This revealed that SST^cs^ animals had a lower PPR than both VIP^cs^ (p<0.01) and WT animals (p<0.01) during CP (**Figure S2A**,**B**). But, during pre-AW, VIP^cs^had a lower PPR over the shorter ISIs compared to controls (**Figure S2B**; p<0.01 for 0.1s and 0.25s ISI). All three backgrounds showed a similar PPR profile during AW (**Figure S2A**,**B**). Our subsequent analysis of MUA responses (**Figure 6E, F**) revealed that silencing SST+ INs abolished the positive PPR for 0.50 ISI (**Figure 6E, G**) observed in controls (**Figure 2E, Figure S2C**). Silencing VIP+ INs resulted in greater adaptation following paired stimuli at short ISIs up until AW (**Figure S2D)**, but otherwise the profile was similar to control animals (**Figure 6H**). These data suggest that GABAergic signalling via SST+ INs play a role in paired-pulse facilitation of MUA observed during CP, whereas an absence of signalling from VIP+ INs impairs the response to stimuli presented at short ISIs, prior to AW.

**Figure 6.**
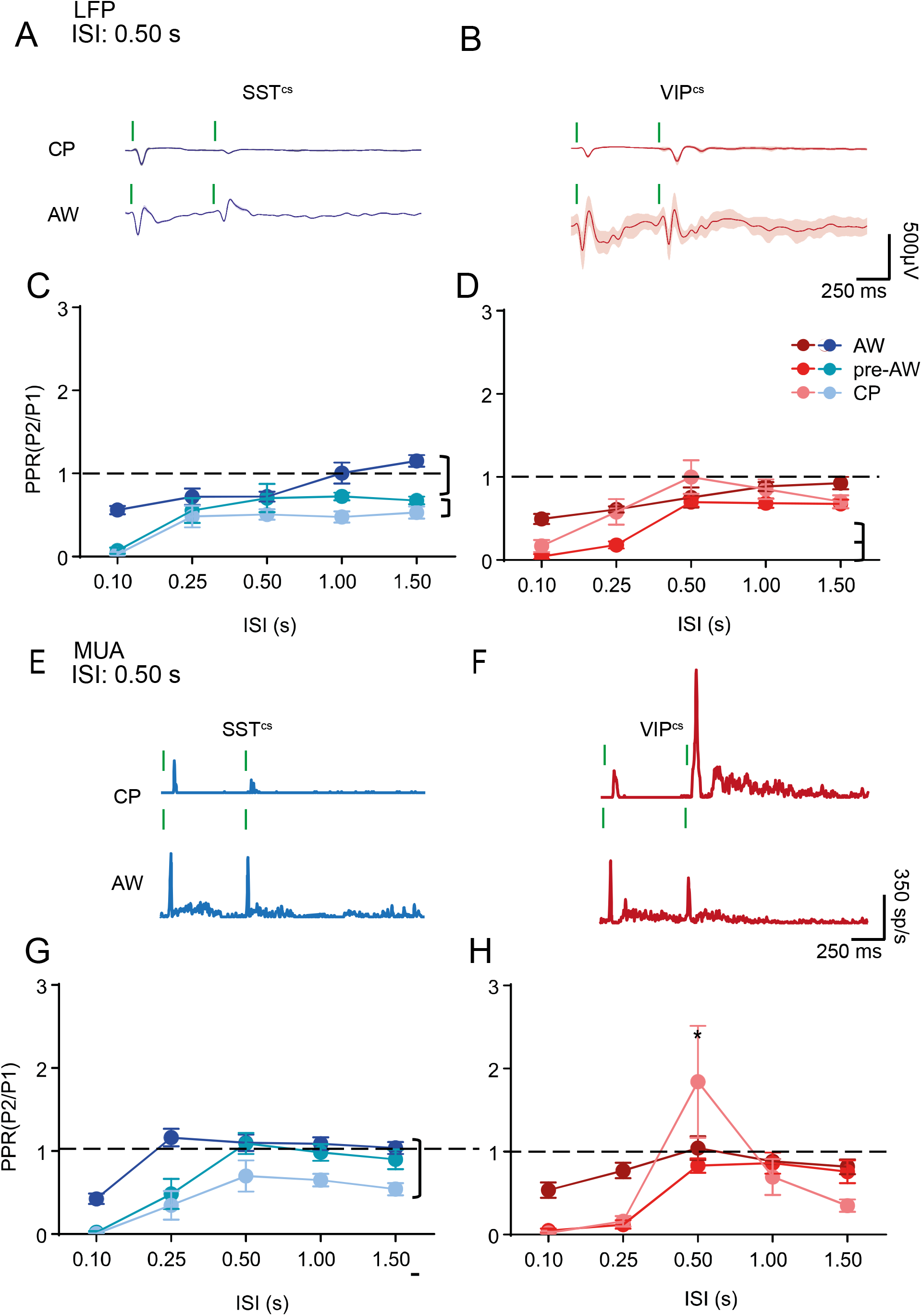
Age-dependent effect of SST+ and VIP+ IN silencing on adaptation. **A**. LFP adaptation response of a SST^cs^ animal to two consecutive deflections with a 0.50s ISI during CP and AW. **B**. LFP adaptation response of VIP^cs^ animal to two consecutive deflections with a 0.50s ISI during CP and AW. **C**. Average PPR of the LFP response of SST^cs^ animals for different ISIs. There was a significant change with ISI(F(4,95) = 18.31, p<0.01) and genotype(F(2,95)= 31.20, p<0.01), but no interaction (F(8,95) = 1.67, p =0.11); CP: N = 9, PAW: N = 6, AW: N =7. **D**. Average PPR of the LFP response of VIP^cs^ animals for different ISIs. There was a significant effect for both ISI (F(4,80) = 20.67, p<0.01) and age (F(2,80) = 10.10, p<0.01) but no interaction(F(8,80) = 1.66, p = 0.12); CP: N = 7, PAW: N = 6, AW: N = 6. **E**. MUA adaptation response of SST^cs^ animal to two consecutive deflections with a 0.50s ISI during CP and AW. **F**. MUA adaptation response of VIP^cs^ animal to two consecutive deflections with a 0.50s ISI during CP and AW. **G**. Average PPR of the MUA response of SST^cs^ animals for different ISIs, There was a significant change with ISI(F(4,95) = 21.59, p<0.01) and genotype(F(2,95)= 27.40, p<0.01), but no interaction (F(8,95) = 1.36, p =0.23); CP: N = 8, PAW: N = 4, AW: N =6. **H**. Average PPR of the MUA response of VIP^cs^ animals for different ISIs. There was an interaction between age and ISI(F(8,50) = 4.10, p < 0.01); CP: N = 4, PAW: N = 6, AW: N = 6. Brackets signify p<0.05 in a simple comparison post ANOVA. * signify p<0.05 in a simple multiple comparison.

### Layer-specific developmental effects of interneuron silencing

GABAergic INs, notably SST+ and VIP+ subtypes, are not evenly distributed across the layers of neocortex (X. Xu, Roby, & Callaway, 2010). Moreover, *in vitro* data has identified the presence of transient translaminar networks mediated by both subtypes during early postnatal life (Marques-Smith et al., 2016; Vagnoni et al., 2020). During the CP time window, MUA is largely confined to granular L4. As such we focused our investigation across supragranular (SG) and infragranular (IG) layers to the pre-AW and AW time windows. During pre-AW, the peak firing rate in SG layers was not affected by silencing of either IN subtype (**Figure 7A**). However we did observed a decrease in IG layer MUA in both VIP^cs^ and SST^cs^ compared to WT animals (p<0.01 in both cases) (**Figure 7C**). The latency of the sensory response was slower across both SG and IG layers in both SST^cs^ and VIP^cs^ animals (**Figure S3A**,**S3C**; p<0.05) compared to wild-type. During AW, both SG and IG response amplitudes of the SST^cs^ animals had increased MUA compared to both VIP^cs^ and WT animals (**Figure 7B;** p<0.05), consistent with an inhibitory role for this IN subtype (Naka et al., 2019). In contrast, the IG response of VIP^cs^animals was lower than WT animals (p<0.05), consistent with a dis-inhibitory role for this IN subtype in IG layers (Pfeffer, Xue, He, Huang, & Scanziani, 2013; Pi et al., 2013). Latencies were similar regardless of genotype (**Figure S3B**,**D**) with the exception of the response latency of SST^cs^ animals at the slowest deflection speed (**Figure S3B**; p<0.05) in the SG layers. The paired-pulse response was consistently altered across layers with responses in both SST^cs^ and VIP^cs^ animals, having lower PPRs, signifying stronger adaptation, than WTs, especially at shorter ISIs (**Figure S4A**,**B;** p<0.05). During the AW time window the PPR of the responses were comparable to controls in line with observations from granular L4 (**Figure S4C**,**D**).

**Figure 7.**
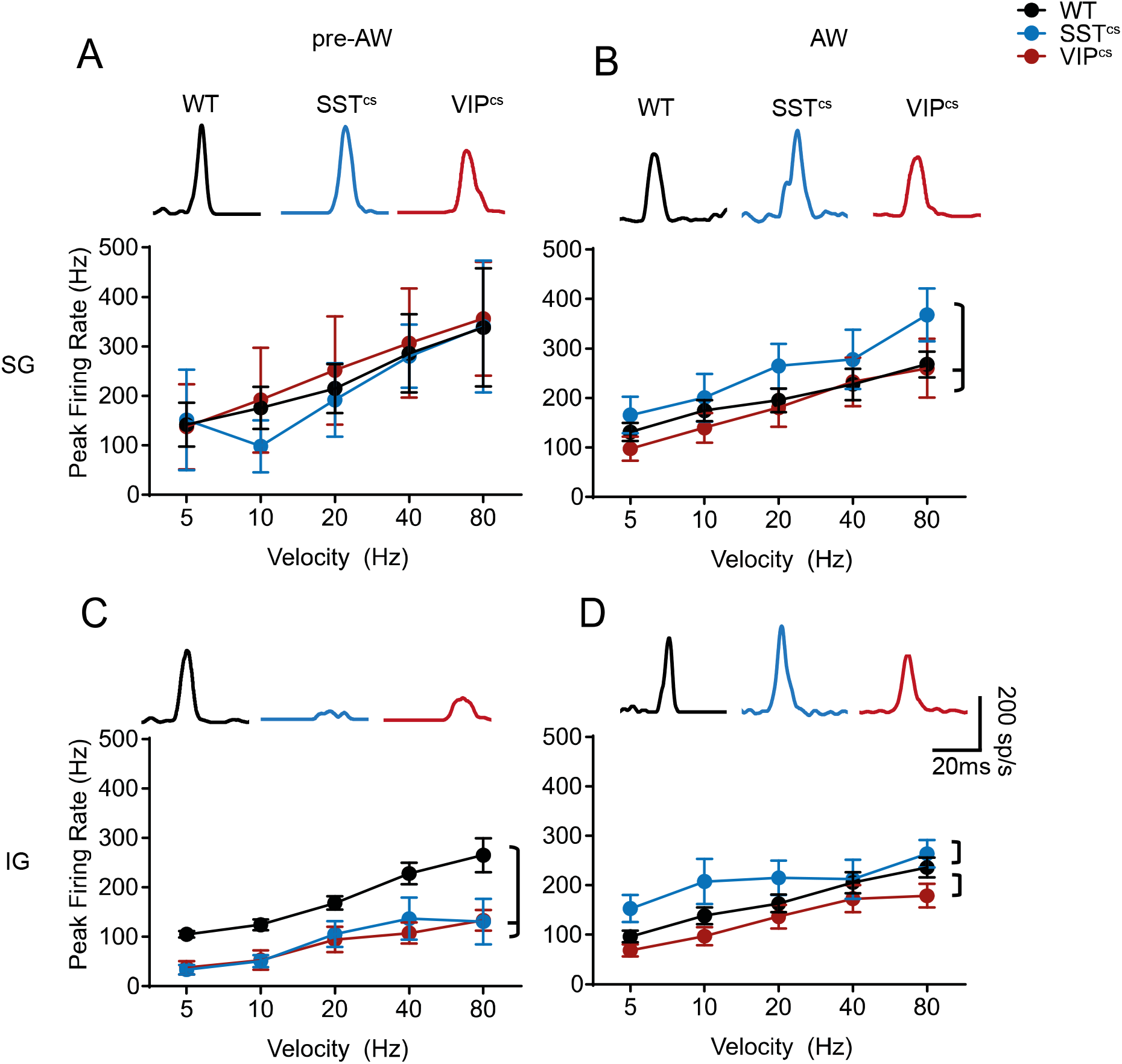
Layer-specific effects of SST+ and VIP+ IN silencing on SER. **A**. Top. SG evoked response traces for the three genotypes. Bottom. The average pre-AW MUA response for a single whisker deflection in the SG layers of WT, SST^cs^ and VIP^cs^ animals. There was an effect for speed(F(4,58) =2.44, p<0.05) but not for genotype(F(2,58) =0.18, p= 0.84 ; WT: N =5, SST^cs^: N =4, VIP^cs^: N= 6) **B**. The average AW MUA response for a single whisker deflection in the SG layers of WT, SST^cs^ and VIP^cs^ animals. There was an effect for both speed (F(4,142) = 7.42, p<0.01) and genotype (F(2,142) = 5.60, p<0.01; WT :N=20, SST^cs^: N = 6 ;VIP^cs^ :N=6) **C**. The average pre-AW MUA response for a single whisker deflection in the IG layers of WT, SST^cs^ and VIP^cs^ animals. There was an effect for both speed(F(4,68) = 13.84, p<0.01) and genotype(F(2,68) = 25.24, p<0.01; WT: N =6, SST^cs^: N =5, VIP^cs^: N= 6) **D**. The average AW MUA response for a single whisker deflection in the IG layers of WT, SST^cs^ and VIP^cs^ animals. There was an effect for both speed (F(4,142) =7.87 ,p<0.01) and genotype(F(2,142) =7.40, p<0.01; WT: N = 21, SST^cs^: N =6, VIP^cs^: N= 6). Brackets signify p<0.05 in a post-hoc multiple comparison.

## Discussion

The contribution of GABAergic interneuron subtypes to early sensory-evoked activity on the millisecond timescale is poorly understood. In this study we have used genetic silencing of GABA release in interneurons to determine the role that SST+ and VIP+ INs play in the acquisition of normal sensory function in S1BF druing the first few weeks of development. Analysis of our *in vivo* data reveals a different role for these two subtypes in mouse: SST+ INs contribute to thalamocortical maturation and plasticity in line with previous *in vitro* circuit analysis, whereas VIP+ INs regulate spiking activity through both inhibitory and dis-inhibitory mechanisms towards the onset of active whisking.

To study the early role of INs we conditionally deleted exons 5 and 6 of the t-SNARE protein Snap25 in SST+ and VIP+ neurons, thereby preventing action potential-dependent neurotransmitter release (Marques-Smith et al., 2016; Verhage et al., 2000; Washbourne et al., 2002). We favoured this approach over alternative genetic manipulations – including expression of potassium rectifier channels – as we felt it was more selective in preventing GABAergic signalling through the time window of our analysis from the onset of the critical period of plasticity to active whisking. We focused on two of the major cortical IN subtypes, SST+ and VIP+ INs, which we could reliably target genetically using Cre lines (Taniguchi et al., 2011). The lack of a specific Cre line for fast spiking, PV+ basket cells at early postnatal ages precluded assessment of these INs during the time window analysed.

Given the key role of locally projecting GABAergic INs in balancing excitation and inhibition in adult neocortex, it is unsurprising that silencing INs led to an increase in spontaneous activity at the later ages tested, at the onset of active perception. However, in the case of VIP+ INs this is counterintuitive given the body of data that suggest these cells exert a primarily dis-inhibitory effect on pyramidal cells via the inhibition of SST+ INs (Pfeffer et al., 2013; Pi et al., 2013). That said, this observation echoes recent findings that suggest that VIP+ INs directly inhibit pyramidal cells in a state-dependent manner (Batista-Brito et al., 2017; Vagnoni et al., 2020). Moreover, it is clear from the effect of silencing INs on spontaneous activity that VIP+ INs develop to control spontaneous pyramidal cell activity around the onset of active whisking. In contrast, our evidence shows that SST+ INs contribute to activity as early as the critical period of plasticity. Silencing action potential-dependent release of GABA from this IN subtype led to a decrease in spontaneous SB and associated spike activity at early ages. Early SB activity is thought to be important for normal sensory development and play a role in the prevention of activity-dependent apoptosis, amongst other formative processes (Khazipov & Milh, 2017). A number of potential mechanisms could explain the effect of SST+ IN silencing on SB generation: first, we broadly targeted SST+ INs. This will affect signalling elsewhere in forebrain, notably the ventral posteromedial nucleus (VPM) of the thalamus. Second, silencing SST+ INs could lead to an increase in local GABAergic signalling through dis-inhibition. Third, silencing SST+ INs could result in delayed maturation of thalamic innervation of neocortex given the role of these cells in early thalamocortical circuits (Yang et al., 2013). The first option is unlikely given that intra-thalamic inhibiton has not fully matured at this age (Zolnik & Connors, 2016) and intra-thalamic SST+ neurons do not synapse onto VPM relay neurons (Clemente-Perez et al., 2017). Our *in vivo* observations are consistent with our previous study that identified both delayed thalamocortical innervation and compensatory increase in the local GABAergic innervation – most likely from immature basket cells (Daw et al., 2007), following SST+ IN silencing *in vitro* (Marques-Smith et al., 2016). Moreover our data are consistent with the thalamic origin of SB generation (Marat Minlebaev et al., 2007) but further support the notion that interneurons circuits across infra- and granular layers interpret afferent sensory signalling to constrain and direct circuit development (Marques-Smith et al., 2016).

Encoding velocity is a key requirement for somatosensory detection by the vibrissae (Kleinfeld et al., 2006; Szwed, Bagdasarian, & Ahissar, 2003). We could detect speed coding in the cortex from the earliest postnatal time point recorded. This suggests that while this sensory computation develops independent of cortical maturation, probably as a result of phase coding as early as at the brainstem level of sensory processing (Szwed et al., 2003; Wallach, Bagdasarian, & Ahissar, 2016). This is entirely consistent with other reports that have identified various stimulus properties encoded in the VPM in adult animals, including speed (Bale, Ince, Santagata, & Petersen, 2015). Further support for upstream processing of speed comes from the lack of an effect for SST+ or VIP+ IN silencing on this computation.

In contrast to speed coding, the profile of sensory response adaptation changed over development. In young animals – during the critical period for plasticity in L4, adaptation took an ‘Inverse-U’ shape with significant depression of the second response at both short and long inter-stimulus intervals, while an interval of 0.5s led to its facilitation. The depression of the second response at short intervals in young animals can likely be explained by low release probability of the immature thalamocortical synapses (Crair & Malenka, 1995; Isaac, Crair, Nicoll, & Malenka, 1997). However, this is less likely to underpin the attenuation observed at longer intervals, which could involve recurrent GABAergic networks. Certainly, it would appear that SST+ INs contribute to the observed facilitation at 0.5s interval as this is abolished in animals in which SST INs are silenced. This can be either directly through facilitation of the TC input onto pyramidal cells through excitatory GABAergic signalling (Ben-Ari, 2002), or indirectly through SST+ IN influence on local GABAergic circuits, as was shown before in the infragranular layers of S1BF (Tuncdemir et al., 2016). Another possibility is that the altered facilitation in SST^CS^ animals is a by-product of the attenuated thalamic input in these animals. However, it is less likely, as we do not observe any impact of VIP+ IN silencing on the facilitation despite these animals also exhibiting reduced thalamic input. Of note is the fact that the 0.5s interval corresponds to the frequency of whisker stimulation that evokes the largest haemodynamic response in S1BF (Sintsov, Suchkov, Khazipov, & Minlebaev, 2017), and results in long term potentiation (LTP) during this developmental time window (An, Yang, Sun, Kilb, & Luhmann, 2012). Together these lines of evidence support the notion that this specific interval is physiologically relevant for the young animal, matching slow passive stimuli it receives, mostly from its littermates (Akhmetshina, Nasretdinov, Zakharov, Valeeva, & Khazipov, 2016). Moreover that it requires functional SST+ IN network.

In adult mice, SST+ and VIP+ INs have been shown to have layer-specific functions (Muñoz et al., 2017; Pfeffer et al., 2013; Pi et al., 2013; H. Xu, Jeong, Tremblay, & Rudy, 2013). Though our work focused primarily on granular L4 – given the consistency of the sensory-evoked response in this layer, we did observe layer-specific changes in sensory processing in our genetically modified mice across the developmental time window tested. In adult mice SST+ INs exert an inhibitory effect in the upper layers but are dis-inhibitory in L4 (H. Xu et al., 2013). We observed an increase in sensory-evoked spiking in supragranular layers after the onset of active whisking, suggesting that the inhibitory effect reported in adults emerges in line with active somato-sensation. VIP+ INs, present mostly in upper layers, have a dis-inhibitory role in the mature cortex (Muñoz et al., 2017; Pfeffer et al., 2013; Pi et al., 2013). In our animals, silencing this IN subtype led to a decrease in response, consistent with dis-inhibition. However, this effect was mostly in the lower, infragranular layers. We did not observe any change in supragranular layer activity, in line with the late integration of VIP+ INs in the local network (Batista-Brito et al., 2017; Vagnoni et al., 2020). Before the onset of active sensation, VIP+ IN silencing led to a transient increase in sensory response in the granular layer. This is consistent with previous findings showing an increase in synapses between these INs and pyramidal cells during this time period (Vagnoni et al., 2020). Overall, the changes we observed are consistent with an inside-out pattern of innervation involving both IN subtypes, whereby the interneurons first integrate in infragranular layers before sequentially innervating supragranular target neurons.

Taken together, our results show that sensory processing develops in line with cortical maturation. We demonstrate that SST+ and VIP+ INs both contribute to early processing of sensory information, with SST+ INs having a distinct role in early regulation of spontaneous activity and facilitation. VIP+ INs play more of a role toward the onset of active perception, regulating incoming sensory information. Our data identify the importance of IN diversity in *in vivo* cortical processing, across early postnatal development.

## Author Contribution

LJB, MK and SJBB conceived experiments which were conducted by LJB, who in addition analysed the data. LJB and SJBB wrote the manuscript. All authors edited the manuscript.

## Acknowledgments

Research in the lab was funded by a BBSRC project grant (BB/P003796/1) and a Medical Sciences Internal Fund: Pump Priming grant (0006784) awarded to LJB. We would also like to thank the Micron Advanced Bioimaging Unit (supported by Wellcome Strategic Awards 736 091911/B/10/Z and 107457/Z/15/Z) for their assistance in this work.

## Supplementary Figures

**Figure S1.**
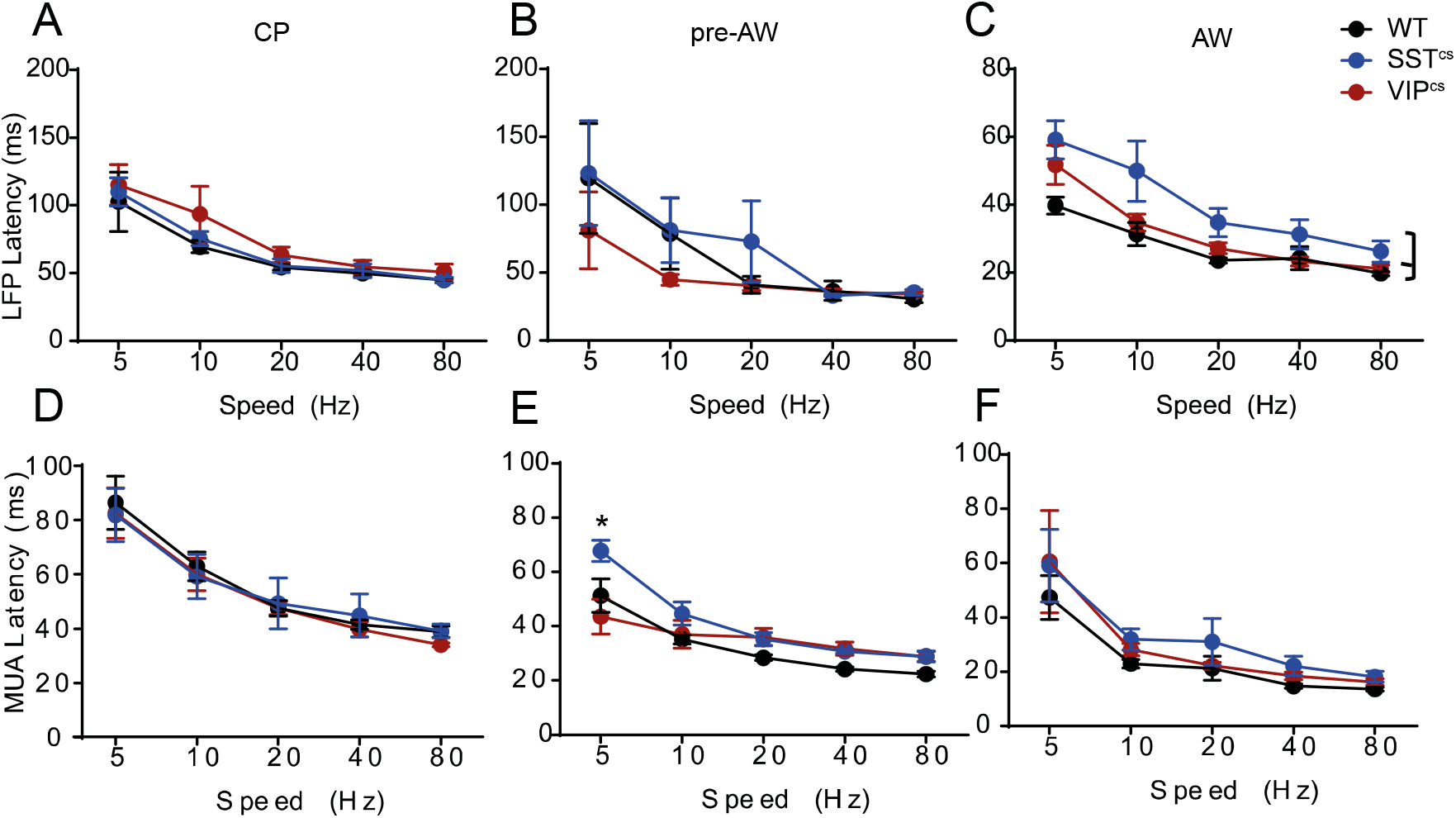
**A**. The average peak LFP latency of WT, SST^cs^ and VIP^cs^ animals during CP. There was a significant effects for speed (F(4,98) = 25.26, p<0.001) but not for genotype(F(2,98) = 2.14, p=0.123). **B**. The average LFP latency of WT, SST^cs^ and VIP^cs^ animals during pre-AW. There was a significant effects for speed (F(4,85) = 5.826, p<0.001) but not for genotype (F(2,85) = 1.148, p=0.322). **C**. The average LF latency of WT, SST^cs^ and VIP^cs^ animals during AW. There was a significant effect for speed (F(4,152) = 24.33, p<0.001) and for genotype(F(2,152) = 15.35, p<0.001). **D**. The MUA latency of WT, SST^cs^ and VIPcs animals during CP. There was an effect for speed (F(4,70) = 30.04, p<0.01), but not for genotype (F(2,70) = 0.75, p=0.475) **E**. The average MUA latency of WT, SST^cs^ and VIP^cs^ animals during pre-AW. There was a genotype X speed interaction (F(8,60) = 2.374, p< 0.05), due to SST^cs^ being slower at 1Hz deflections (p<0.05). **F**. The average MUA latency of WT, SST^cs^ and VIP^cs^ animals during AW. There was an effect for speed (F(4,147) = 15.29, p<0.001), but not for genotype (F(2,147) = 2.488, p = 0.086). Brackets signify p<0.05 for a post-hoc multiple comparison. * p<0.05 for a simple post-hoc multiple comparison.

**Figure S2.**
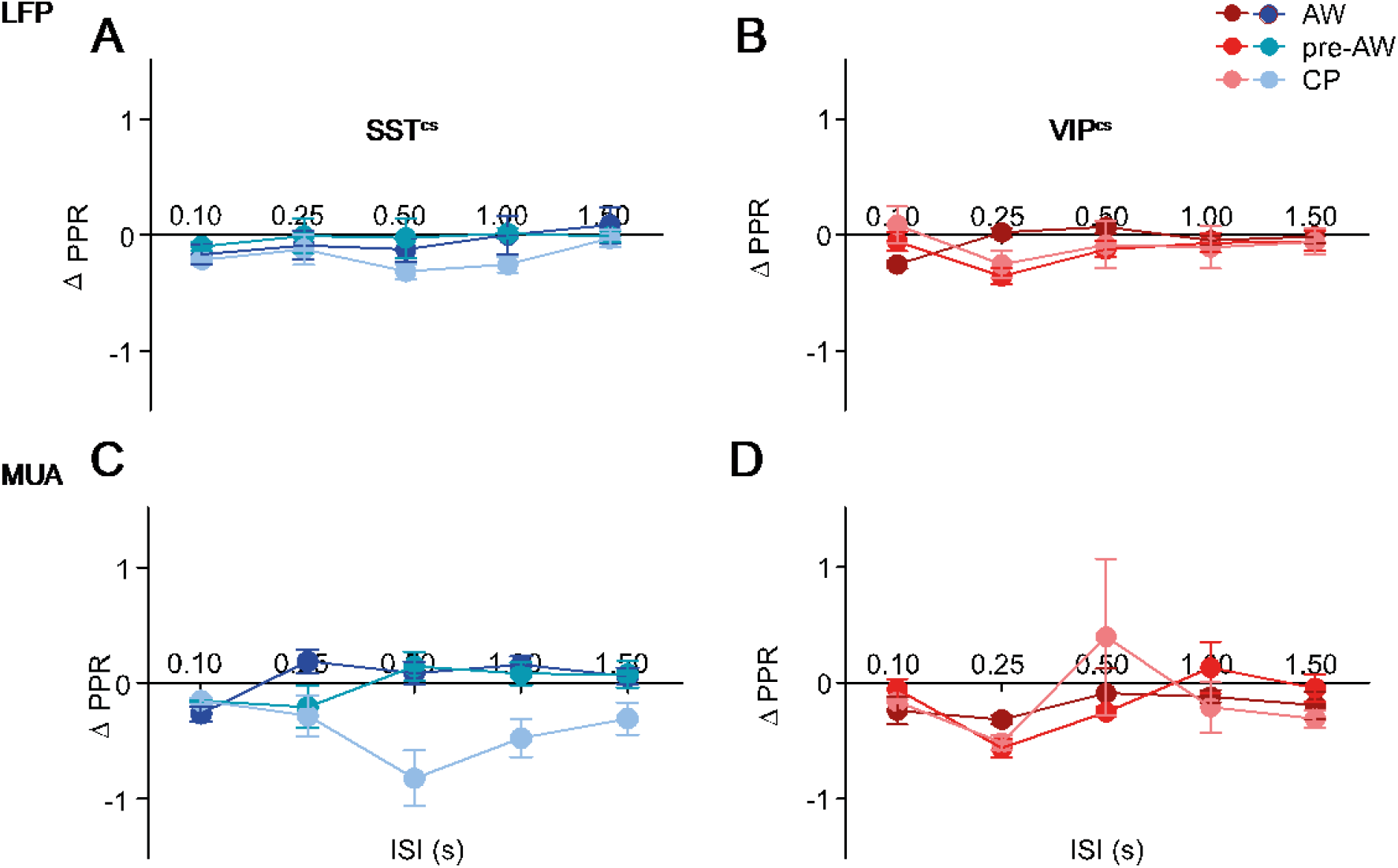
**A**. The average difference in LFP PPR of SST^cs^ animals compared to the average in controls over development. **B**. The average difference in LFP PPR of VIP^cs^ animals compared to the average in controls over development. **C**. The average difference in MUA PPR of SST^cs^ animals compared to the average in controls over development. **D**. The average difference in MUA PPR of VIP^cs^animals compared to the average in controls over development.

**Figure S3.**
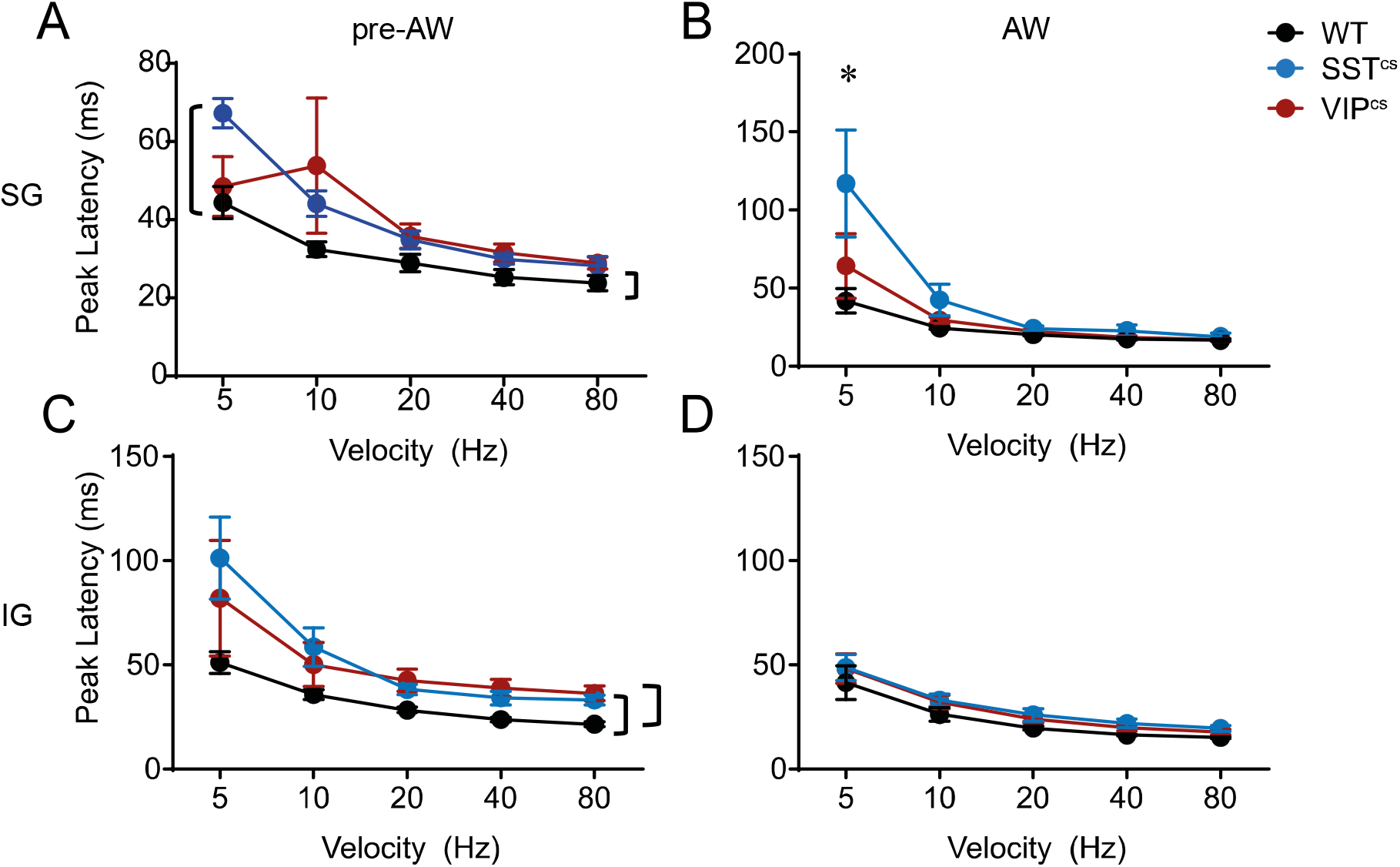
The effect of VIP+ and SST+ IN silencing of response latency. **A**. Average response latencies in SG layers during pre-AW. There was an effect for both speed(F(4,58) =11.51, p<0.01) and genotype(F(2,58) =4.71, p<0.01). **B**. Average response latencies in SG layers during AW. There was an interaction between genotype and speed (F(8,142) = 4.44, p<0.01). The interaction stemmed from the 5 Hz deflection where both silenced genotypes had slower latencies (p<0.01). **C**. average response latencies in IG layers during pre-AW. There was an effect for both speed(F(4,68) = 14.10, p<0.01) and genotype(F(2,68) = 8.17, p<0.01). **D**. Average response latencies in IG layers during AW. There was only an effect for speed (F(4,142) = 13.22, p<0.01), but not for genotype (F(2,142) = 2.31, p=0.10)

**Figure S4.**
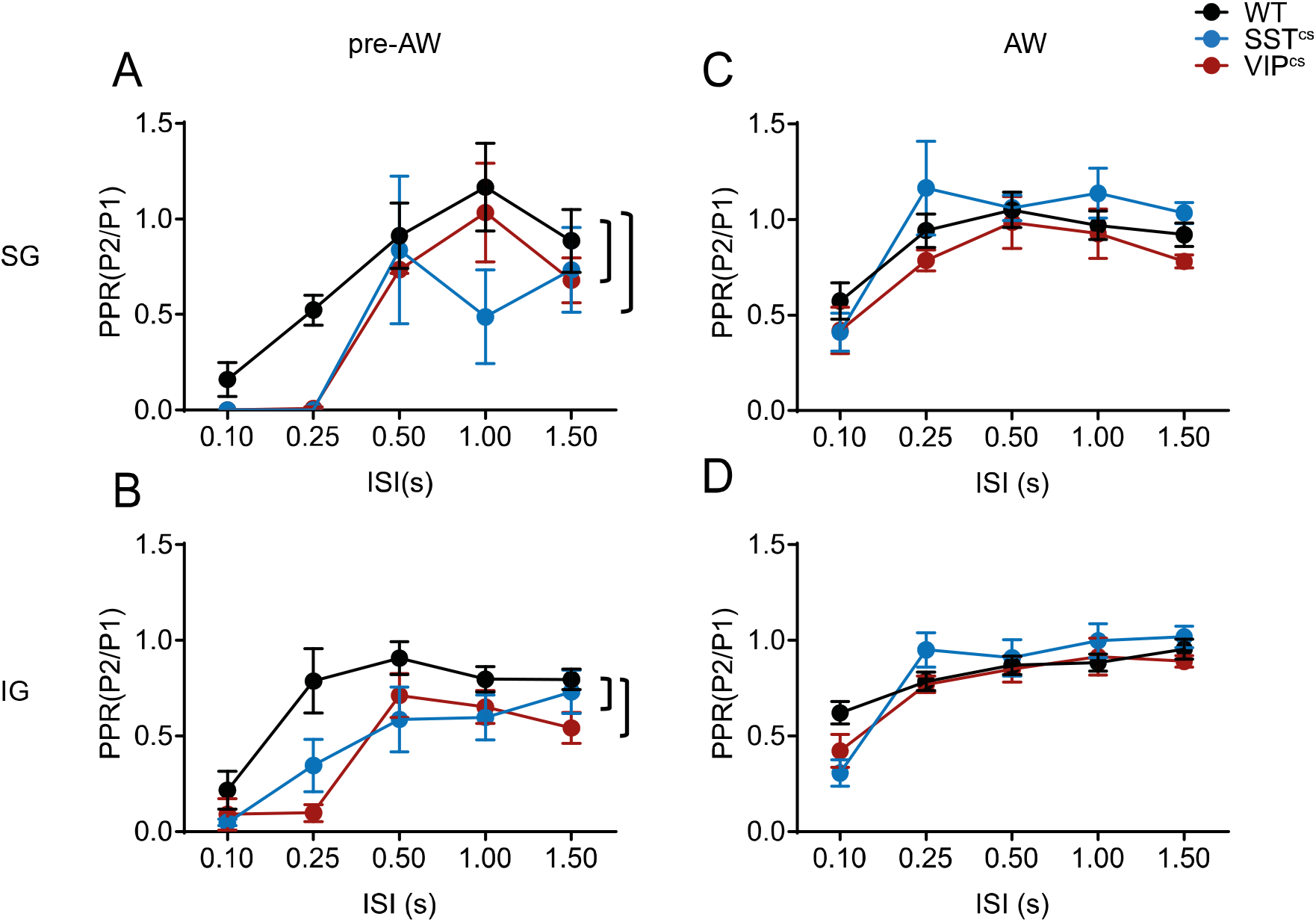
**A**. The average PPR of the MUA response in the SG layer of WT, SST^cs^, and VIP^cs^animals during pre-AW. There was an effect for both ISI (F(4,100) = 11.61, p<0.01) and genotype (F(2,100) = 4.336, p<0.05; WT: N = 13, SST^cs^: N = 4, VIP^cs^: N = 6). **B**. The average PPR of the MUA response in the IG layer of WT, SST^cs^, and VIP^cs^ animals during pre-AW. There was an effect for both ISI (F(4,120) = 11.40, p<0.01) and genotype (F(2,120) = 9.013, p<0.01; WT: N = 18, SST^cs^: N = 4, VIP^cs^: N = 6). **C**. The average PPR of the MUA response in the SG layer of WT, SST^cs^, and VIP^cs^ animals during AW. There was an effect for ISI (F(4,173) = 8.521, p<0.01), but not for genotype (F(2,173) = 1.673, p=0.191; WT: N = 27, SST^cs^: N = 5, VIP^cs^: N = 6) **D**. The average PPR of the MUA response in the IG layer of WT, SST^cs^, and VIP^cs^ animals during AW. There was an effect for ISI (F(4,188) = 16.46, p<0.01) but not for genotype (F(2,88 = 0.671, p=0.512; WT: N = 30, SST^cs^: N = 6, VIP^cs^: N = 6)

## References

Akhmetshina, D., Nasretdinov, A., Zakharov, A., Valeeva, G., & Khazipov, R. (2016). The nature of the sensory input to the neonatal rat barrel cortex. Journal of Neuroscience. https://doi.org/10.1523/JNEUROSCI.1781-16.2016

Allitt, B. J., Alwis, D. S., & Rajan, R. (2017). Laminar-specific encoding of texture elements in rat barrel cortex. Journal of Physiology. https://doi.org/10.1113/JP274865

An, S., Yang, J. W., Sun, H., Kilb, W., & Luhmann, H. J. (2012). Long-term potentiation in the neonatal rat barrel cortex in vivo. Journal of Neuroscience. https://doi.org/10.1523/JNEUROSCI.1212-12.2012

Anastasiades, P. G., & Butt, S. J. B. (2012). A role for silent synapses in the development of the pathway from layer 2/3 to 5 pyramidal cells in the neocortex. Journal of Neuroscience. https://doi.org/10.1523/JNEUROSCI.1262-12.2012

Anastasiades, P. G., Marques-Smith, A., Lyngholm, D., Lickiss, T., Raffiq, S., Katzel, D., … Butt, S. J. B. (2016). GABAergic interneurons form transient layer-specific circuits in early postnatal neocortex. Nature Communications. https://doi.org/10.1038/ncomms10584

Arabzadeh, E., Petersen, R. S., & Diamond, M. E. (2003). Encoding of whisker vibration by rat barrel cortex neurons: Implications for texture discrimination. Journal of Neuroscience. https://doi.org/10.1523/jneurosci.23-27-09146.2003

Ayzenshtat, I., Karnani, M. M., Jackson, J., & Yuste, R. (2016). Cortical control of spatial resolution by VIP+ interneurons. Journal of Neuroscience. https://doi.org/10.1523/JNEUROSCI.1920-16.2016

Bale, M. R., Ince, R. A. A., Santagata, G., & Petersen, R. S. (2015). Efficient population coding of naturalistic whisker motion in the ventro-posterior medial thalamus based on precise spike timing. Frontiers in Neural Circuits. https://doi.org/10.3389/fncir.2015.00050

Batista-Brito, R., Vinck, M., Ferguson, K. A., Chang, J. T., Laubender, D., Lur, G., … Cardin, J. A. (2017). Developmental Dysfunction of VIP Interneurons Impairs Cortical Circuits. Neuron, 95(4), 884-895.e9. https://doi.org/10.1016/J.NEURON.2017.07.034

Ben-Ari, Y. (2002). Excitatory actions of GABA during development: The nature of the nurture. Nature Reviews Neuroscience. https://doi.org/10.1038/nrn920

Ben-Ari, Y., Khalilov, I., Represa, A., & Gozlan, H. (2004). Interneurons set the tune of developing networks. Trends in Neurosciences, Vol. 27, pp. 422–427. https://doi.org/10.1016/j.tins.2004.05.002

Butt, S., Fuccillo, M., Nery, S., Noctor, S., Kriegstein, A., Corbin, J. G., & Fishell, G. (2005). The temporal and spatial origins of cortical interneurons predict their physiological subtype. Neuron, 48(4), 591–604. https://doi.org/10.1016/j.neuron.2005.09.034

Butt, S. J., Stacey, J. A., Teramoto, Y., & Vagnoni, C. (2017). A role for GABAergic interneuron diversity in circuit development and plasticity of the neonatal cerebral cortex. Current Opinion in Neurobiology. https://doi.org/10.1016/j.conb.2017.03.011

Carvell, G. E., & Simons, D. J. (1990). Biometric analyses of vibrissal tactile discrimination in the rat. Journal of Neuroscience. https://doi.org/10.1523/jneurosci.10-08-02638.1990

Clemente-Perez, A., Makinson, S. R., Higashikubo, B., Brovarney, S., Cho, F. S., Urry, A., … Paz, J. T. (2017). Distinct Thalamic Reticular Cell Types Differentially Modulate Normal and Pathological Cortical Rhythms. Cell Reports. https://doi.org/10.1016/j.celrep.2017.05.044

Crair, M. C., & Malenka, R. C. (1995). A critical period for long-term potentiation at thalamocortical synapses. Nature. https://doi.org/10.1038/375325a0

Daw, M. I., Ashby, M. C., & Isaac, J. T. R. (2007). Coordinated developmental recruitment of latent fast spiking interneurons in layer IV barrel cortex. Nature Neuroscience, 10(4), 453–461. https://doi.org/10.1038/nn1866

Doischer, D., Hosp, J. A., Yanagawa, Y., Obata, K., Jonas, P., Vida, I., & Bartos, M. (2008). Postnatal differentiation of basket cells from slow to fast signaling devices. Journal of Neuroscience. https://doi.org/10.1523/JNEUROSCI.2890-08.2008

Erzurumlu, R. S., & Gaspar, P. (2012). Development and critical period plasticity of the barrel cortex. European Journal of Neuroscience. https://doi.org/10.1111/j.1460-9568.2012.08075.x

Guić-Robles, E., Valdivieso, C., & Guajardo, G. (1989). Rats can learn a roughness discrimination using only their vibrissal system. Behavioural Brain Research. https://doi.org/10.1016/0166-4328(89)90011-9

Hanganu-Opatz, I. L., Butt, S. J. B., Hippenmeyer, S., De Marco García, N. V., Cardin, J. A., Voytek, B., & Muotri, A. R. (2021). The Logic of Developing Neocortical Circuits in Health and Disease. The Journal of Neuroscience. https://doi.org/10.1523/jneurosci.1655-20.2020

Hensch, T. K. (2005). Critical period plasticity in local cortical circuits. Nature Reviews Neuroscience. https://doi.org/10.1038/nrn1787

Isaac, J. T. R., Crair, M. C., Nicoll, R. A., & Malenka, R. C. (1997). Silent synapses during development of thalamocortical inputs. Neuron. https://doi.org/10.1016/S0896-6273(00)80267-6

Khazipov, Roustem, & Milh, M. (2017). Early patterns of activity in the developing cortex: Focus on the sensorimotor system. Seminars in Cell & Developmental Biology. https://doi.org/10.1016/J.SEMCDB.2017.09.014

Khazipov, Rustom, Sirota, A., Leinekugel, X., Holmes, G. L., Ben-Ari, Y., & Buzsáki, G. (2004). Early motor activity drives spindle bursts in the developing somatosensory cortex. Nature, 432(7018), 758–761. https://doi.org/10.1038/nature03132

Kleinfeld, D., Ahissar, E., & Diamond, M. E. (2006). Active sensation: insights from the rodent vibrissa sensorimotor system. Current Opinion in Neurobiology. https://doi.org/10.1016/j.conb.2006.06.009

Kolasinski, J., Logan, J. P., Hinson, E. L., Manners, D., Divanbeighi Zand, A. P., Makin, T. R., … Stagg, C. J. (2017). A Mechanistic Link from GABA to Cortical Architecture and Perception. Current Biology. https://doi.org/10.1016/j.cub.2017.04.055

Malina, K. C. K., Mohar, B., Rappaport, A. N., & Lampl, I. (2016). Local and thalamic origins of correlated ongoing and sensory-evoked cortical activities. Nature Communications. https://doi.org/10.1038/ncomms12740

Maravall, M., Petersen, R. S., Fairhall, A. L., Arabzadeh, E., & Diamond, M. E. (2007). Shifts in coding properties and maintenance of information transmission during adaptation in barrel cortex. PLoS Biology. https://doi.org/10.1371/journal.pbio.0050019

Marques-Smith, A., Lyngholm, D., Kaufmann, A. K., Stacey, J. A., Hoerder-Suabedissen, A., Becker, E. B. E., … Butt, S. J. B. (2016). A Transient Translaminar GABAergic Interneuron Circuit Connects Thalamocortical Recipient Layers in Neonatal Somatosensory Cortex. Neuron, 89(3), 536–549. https://doi.org/10.1016/j.neuron.2016.01.015

McRae, P. A., Rocco, M. M., Kelly, G., Brumberg, J. C., & Matthews, R. T. (2007). Sensory deprivation alters aggrecan and perineuronal net expression in the mouse barrel cortex. Journal of Neuroscience. https://doi.org/10.1523/JNEUROSCI.5425-06.2007

Minlebaev, M., Colonnese, M., Tsintsadze, T., Sirota, A., & Khazipov, R. (2011). Early Gamma Oscillations Synchronize Developing Thalamus and Cortex. Science, 334(6053), 226–229. https://doi.org/10.1126/science.1210574

Minlebaev, Marat, Ben-Ari, Y., & Khazipov, R. (2007). Network Mechanisms of Spindle-Burst Oscillations in the Neonatal Rat Barrel Cortex In Vivo. Journal of Neurophysiology, 97(1), 692–700. https://doi.org/10.1152/jn.00759.2006

Miyoshi, G., Young, A., Petros, T., Karayannis, T., Chang, M. M. K., Lavado, A., … Fishell, G. (2015). Prox1 regulates the subtype-specific development of caudal ganglionic eminence-derived GABAergic cortical interneurons. Journal of Neuroscience. https://doi.org/10.1523/JNEUROSCI.1164-15.2015

Modol, L., Bollmann, Y., Tressard, T., Baude, A., Che, A., Duan, Z. R. S., … Cossart, R. (2020). Assemblies of Perisomatic GABAergic Neurons in the Developing Barrel Cortex. Neuron. https://doi.org/10.1016/j.neuron.2019.10.007

Muñoz, W., Tremblay, R., Levenstein, D., & Rudy, B. (2017). Layer-specific modulation of neocortical dendritic inhibition during active wakefulness. Science, 355(6328), 954–959. https://doi.org/10.1126/science.aag2599

Musall, S., Haiss, F., Weber, B., & von der Behrens, W. (2017). Deviant Processing in the Primary Somatosensory Cortex. Cerebral Cortex (New York, N.Y.: 1991), 27(1), 863–876. https://doi.org/10.1093/cercor/bhv283

Naka, A., Veit, J., Shababo, B., Chance, R. K., Risso, D., Stafford, D., … Adesnik, H. (2019). Complementary networks of cortical somatostatin interneurons enforce layer specific control. ELife, 8. https://doi.org/10.7554/eLife.43696

Natan, R. G., Briguglio, J. J., Mwilambwe-Tshilobo, L., Jones, S. I., Aizenberg, M., Goldberg, E. M., & Geffen, M. N. (2015). Complementary control of sensory adaptation by two types of cortical interneurons. ELife, 4(OCTOBER 2015). https://doi.org/10.7554/eLife.09868

Nicholson, C., & Freeman, J. A. (1975). Theory of current source density analysis and determination of conductivity tensor for anuran cerebellum. Journal of Neurophysiology. https://doi.org/10.1152/jn.1975.38.2.356

Nowicka, D., Soulsby, S., Skangiel-Kramska, J., & Glazewski, S. (2009). Parvalbumin-containing neurons, perineuronal nets and experience-dependent plasticity in murine barrel cortex. European Journal of Neuroscience. https://doi.org/10.1111/j.1460-9568.2009.06996.x

Oh, W. C., Lutzu, S., Castillo, P. E., & Kwon, H. B. (2016). De novo synaptogenesis induced by GABA in the developing mouse cortex. Science. https://doi.org/10.1126/science.aaf5206

Ollerenshaw, D. R., Zheng, H. J. V., Millard, D. C., Wang, Q., & Stanley, G. B. (2014). The adaptive trade-off between detection and discrimination in cortical representations and behavior. Neuron, 81(5), 1152–1164. https://doi.org/10.1016/j.neuron.2014.01.025

Pachitariu, M., Steinmetz, N., Kadir, S., Carandini, M., & Kenneth D. H. (2016). Kilosort: realtime spike-sorting for extracellular electrophysiology with hundreds of channels. BioRxiv, 061481. https://doi.org/10.1101/061481

Paxinos, G., Halliday, G., Watson, C., & Mustafa, K. (2020). Atlas of the Developing Mouse Brain (2nd ed.). Academic Press.

Petersen, C. C. H. (2007). The functional organization of the barrel cortex. Neuron, 56(2), 339–355. https://doi.org/10.1016/j.neuron.2007.09.017

Petersen, R. S., Panzeri, S., & Diamond, M. E. (2002). Population coding in somatosensory cortex. Current Opinion in Neurobiology. https://doi.org/10.1016/S0959-4388(02)00338-0

Pfeffer, C. K., Xue, M., He, M., Huang, Z. J., & Scanziani, M. (2013). Inhibition of inhibition in visual cortex: The logic of connections between molecularly distinct interneurons. Nature Neuroscience. https://doi.org/10.1038/nn.3446

Pi, H. J., Hangya, B., Kvitsiani, D., Sanders, J. I., Huang, Z. J., & Kepecs, A. (2013). Cortical interneurons that specialize in disinhibitory control. Nature. https://doi.org/10.1038/nature12676

Rappelsberger, P., Pockberger, H., & Petsche, H. (1981). Current source density analysis: Methods and application to simultaneously recorded field potentials of the rabbit’s visual cortex. Pflügers Archiv European Journal of Physiology. https://doi.org/10.1007/BF00582108

Rudy, B., Fishell, G., Lee, S. H., & Hjerling-Leffler, J. (2011). Three groups of interneurons account for nearly 100% of neocortical GABAergic neurons. Developmental Neurobiology. https://doi.org/10.1002/dneu.20853

Seelke, A. M. H., & Blumberg, M. S. (2010). Developmental appearance and disappearance of cortical events and oscillations in infant rats. Brain Research, 1324, 34–42. https://doi.org/10.1016/J.BRAINRES.2010.01.088

Sintsov, M., Suchkov, D., Khazipov, R., & Minlebaev, M. (2017). Developmental changes in sensory-evoked optical intrinsic signals in the rat barrel cortex. Frontiers in Cellular Neuroscience. https://doi.org/10.3389/fncel.2017.00392

Szwed, M., Bagdasarian, K., & Ahissar, E. (2003). Encoding of vibrissal active touch. Neuron. https://doi.org/10.1016/S0896-6273(03)00671-8

Taniguchi, H., He, M., Wu, P., Kim, S., Paik, R., Sugino, K., … Huang, Z. J. (2011). A Resource of Cre Driver Lines for Genetic Targeting of GABAergic Neurons in Cerebral Cortex. Neuron. https://doi.org/10.1016/j.neuron.2011.07.026

Tuncdemir, S. N., Wamsley, B., Stam, F. J., Osakada, F., Goulding, M., Callaway, E. M., … Fishell, G. (2016). Early Somatostatin Interneuron Connectivity Mediates the Maturation of Deep Layer Cortical Circuits. Neuron, 89(3), 521–535. https://doi.org/10.1016/j.neuron.2015.11.020

Urban-Ciecko, J., Fanselow, E. E., & Barth, A. L. (2015). Neocortical somatostatin neurons reversibly silence excitatory transmission via GABAb receptors. Current Biology. https://doi.org/10.1016/j.cub.2015.01.035

Vagnoni, C., Baruchin, L. J., Ghezzi, F., Ratti, S., Molnar, Z., & Butt, S. (2020). Ontogeny of the VIP+ interneuron sensory-motor circuit prior to active whisking. BioRxiv, 2020.07.01.182238. https://doi.org/10.1101/2020.07.01.182238

Vaknin, G., DiScenna, P. G., & Teyler, T. J. (1988). A method for calculating current source density (CSD) analysis without resorting to recording sites outside the sampling volume. Journal of Neuroscience Methods. https://doi.org/10.1016/0165-0270(88)90056-8

van der Bourg, A., Yang, J.-W., Reyes-Puerta, V., Laurenczy, B., Wieckhorst, M., Stüttgen, M. C., … Helmchen, F. (2016). Layer-Specific Refinement of Sensory Coding in Developing Mouse Barrel Cortex. Cerebral Cortex, 27(10), 4835– 4850. https://doi.org/10.1093/cercor/bhw280

Verhage, M., Maia, A. S., Plomp, J. J., Brussaard, A. B., Heeroma, J. H., Vermeer, H., … Südhof, T. C. (2000). Synaptic assembly of the brain in the absence of neurotransmitter secretion. Science. https://doi.org/10.1126/science.287.5454.864

Wallach, A., Bagdasarian, K., & Ahissar, E. (2016). On-going computation of whisking phase by mechanoreceptors. Nature Neuroscience. https://doi.org/10.1038/nn.4221

Washbourne, P., Thompson, P. M., Carta, M., Costa, E. T., Mathews, J. R., Lopez-Benditó, G., … Wilson, M. C. (2002). Genetic ablation of the t-SNARE SNAP-25 distinguishes mechanisms of neuroexocytosis. Nature Neuroscience. https://doi.org/10.1038/nn783

Wood, K. C., Blackwell, J. M., & Geffen, M. N. (2017). Cortical inhibitory interneurons control sensory processing. Current Opinion in Neurobiology. https://doi.org/10.1016/j.conb.2017.08.018

Xu, H., Jeong, H. Y., Tremblay, R., & Rudy, B. (2013). Neocortical Somatostatin-Expressing GABAergic Interneurons Disinhibit the Thalamorecipient Layer 4. Neuron, 77(1), 155–167. https://doi.org/10.1016/j.neuron.2012.11.004

Xu, X., Roby, K. D., & Callaway, E. M. (2010). Immunochemical characterization of inhibitory mouse cortical neurons: Three chemically distinct classes of inhibitory cells. Journal of Comparative Neurology. https://doi.org/10.1002/cne.22229

Yang, J. W., An, S., Sun, J. J., Reyes-Puerta, V., Kindler, J., Berger, T., … Luhmann, H. J. (2013). Thalamic network oscillations synchronize ontogenetic columns in the newborn rat barrel cortex. Cerebral Cortex, 23(6), 1299–1316. https://doi.org/10.1093/cercor/bhs103

Yu, J., Hu, H., Agmon, A., & Svoboda, K. (2019). Recruitment of GABAergic Interneurons in the Barrel Cortex during Active Tactile Behavior. Neuron. https://doi.org/10.1016/j.neuron.2019.07.027

Zolnik, T. A., & Connors, B. W. (2016). Electrical synapses and the development of inhibitory circuits in the thalamus. Journal of Physiology. https://doi.org/10.1113/JP271880

